# An engineered electroosmotic flow transports unravelled proteins across nanopores

**DOI:** 10.1101/2023.02.17.528930

**Authors:** Adina Sauciuc, Blasco Morozzo, Matthijs Tadema, Mauro Chinappi, Giovanni Maglia

## Abstract

The development of a technology capable of sequencing single proteins holds promise to unravel new biological information hidden in ensemble analysis. However, new techniques must be first developed. In one approach, proteins are unfolded and translocate across a nanopore under an external bias. Unlike DNA, however, proteins do not have a uniform charge, and the electrophoretic force cannot be used to translocate proteins. Here, we show that by introducing sets of charges spaced by ~1 nm an otherwise neutral nanopore an electroosmotic force is created that induces the unidirectional transport of polypeptides, even against relatively strong electrophoretic forces. Unstructured polypeptides and native proteins unfolded with urea produce current signatures as they traversed the nanopore, which could lead to quick protein identifcation. This approach can be used to translocate and stretch proteins in non-enzymatic protein identification and enzymatic protein sequencing approaches.

## Introduction

Fast, cheap and high-throughput sequencing of nucleic acids has revolutionised medicine and our understanding of biology. Similar advances in protein sequencing are promising a similar impact. As already demonstrated for nucleic acids, nanopores might also be used for single-molecule protein identification. However, one of the fundamental problems that has to be solved is to capture and translocate polypeptides, and to linearise them as they traverse the nanopore.

Polymers have been transported across nanopores using the electric field that generates near and inside the nanopore by an external voltage. The resulting electrophoretic force (EPF) on charged molecules than drives the entry, transport and stretch of charged polymers such as nucleic acids across nanopores. Proteins, however, have a heterogeneous charge distribution and this approach cannot be used as a means of transport. In fact, the EPF facilitates the transport of certain amino acids while opposing the transport of others as a polypeptide traverses the nanopore.

One option is to use the electroosmotic flow (EOF) - the net flux of water through the nanopore generated by a net ion-charge accumulation inside the nanopore - as a driving force. However, despite it is known that the electroosmotic flow can facilitate the capture^1–3^ or traping^4–6^ of molecules within nanopores, the resulting electroosmotic force correlates with the size and shape of the molecule, and it has yet to be established to what extent, if at all, it can drive the transport and stretching of polymers across nanopores.

Previous work used combinations of EPF and the EOF to transport linearised peptides and protein across nanopores. For example, peptides were attached to a DNA strand and driven into the nanopore by electrophoresis, and then moved by out of the nanopore using an ATP-fuelled helicase.^7–9^ It was found, however, that the neutral peptides cannot be fully stretched by this approach.^9^ Other examples, proteins were elongated with DNA^10–12^ or negatively charged polypeptide tags,^13,14^ which then produced an electrophoretic drag to capture the polypeptide. Then, enzymes or the combination of the EPF and EOF induced the translocation of polypeptides.

It still remains to be discovered whether proteins can be transported as single-file across nanopores using the EOF alone, a likely prerequisite for enzymatic protein sequencing. Notably, the EOF should be strong, as native proteins occasionally contain stretches of charges that will require transport against a focussed electrophoretic force. In this work we show that an EOF can be engineered to induce the transport of relatively highly charged polypeptides against an EPF. This approach, which does not require the use of denaturant, allows the transport and stretching of proteins in both enzymatic and non-enzymatic approaches to protein sequencing.

## Results

### Engineering the electro-osmotic flow in CytK nanopores

In this work we used CytK nanopores (**Figure** 1A). As shown by homology modelling, the nanopore is formed by a spherical vestibule ~5 nm in diameter connected to a ~5×2 nm cylindrical β-barrel region (**Figure** 1B). The latter dominates the electric resistance of the nanopore. The lumen of the wild-type nanopore has no overall charge (the β-barrel region contains two pairs of opposite charge residues: K128-E139 and K155-E112). Consequently, the ion selectivity [(p(K)/p(Cl)] of WT-CytK is nearly one (0.99±0.079, **Figure** 1C), indicating that the WT pore is non-selective and thus shows no electroosmotic flow.

**Figure 1.**
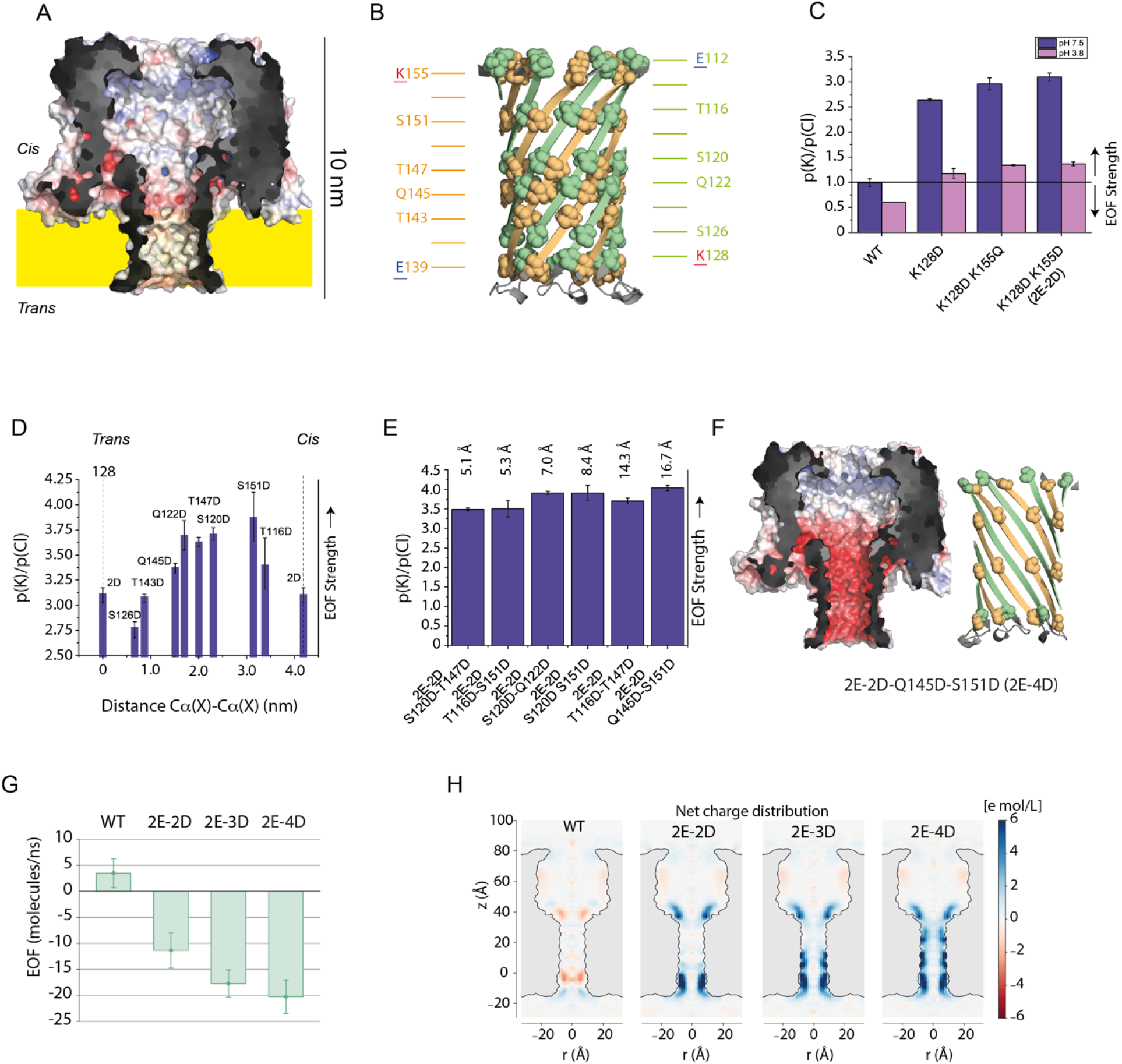
Engineering and electroosmotic flow. **A)** Cut-through of a surface representation of WT-CytK nanopores in 1 M KCl, pH 7.5. The nanopore was made by homology modelling from the αHL nanopore.^19^ **B)** Cartoon representation of WT-CytK β-barrel region. The N-terminal strand is in green and the C-terminal strand in orange. The charged residues are underlined. **C)** Ion selectivity of CytK mutants where the positive charge residues are introduced within the barrel of the nanopore at pH 7.5 (blue) and 3.8 (pink). **D)**Ion selectivity of CytK-2D-2E with an additional engineered aspartate residues at different positions within the β-barrel region. **E)** Ion selectivity for engineered CytK with four additional aspartate residues distributed within the lumen. **F)** Cut-through of a surface (left) and cartoon (right) representation of the most ion selective mutant, CytK-2E-4D, in 1 M KCl, pH 7.5 [(p(K)/p(Cl) of 4.04 ± 0.07, at pH 7.5 and p(K)/p(Cl) of 1.36±0.00 at pH 3.8. **G**) EOF evaluated from MD simulations for WT and different mutants for ΔV = −125 mV. In our reference system, negative EOF corresponds to flows from *cis* to *trans*. EOFs are obtained averaging over 120 ns. **H**) Equilibrium (ΔV = 0) ionic charge distributions. WT is slightly anion selective while K128D-K155D (2E-2D), K128D-K155D-Q145D (2E-3D) and K128D-K155D-S151D (2E-4D) show increasing cation selectivity. Charge density is colour coded with the scale on the right, expressed in molar units. Equilibrium maps are averaged over 40 ns. The cutout of the pore is drawn to lead the eye and it roughly corresponds to the region inaccessible to the solvent.

An EOF was induced in WT-CytK by lowering the pH to 3.8, which in turn increased the overall positive charge of the nanopore by protonation of the acidic residues (E112 and E139) making the nanopore anion selective[(p(K)/p(Cl)= 0.606±0.005]. Alternatively, an electroosmotic flow at physiological pH was introduced by removing a positively charged residues near the *cis* [K128D-CytK, (p(K)/p(Cl) = 2.63±0.02] and *trans* (K128D-K155Q-CytK and K128D-K155D-CytK) entries of the β-barrel region. These latter modifications increased the ion selectivity, but only slightly [(p(K)/p(Cl) = 2.96±0.12 and 3.10±0.08, respectively]. At pH 3.8, the nanopores remained cation selective, but only weakly [(p(K)/p(Cl) = 1.36±0.10 for K128D-K155D-CytK].

To test how much the ion selectivity could be increased in CytK, we introduced an additional negative charge at different positions within the β-barrel of K128D-K155D-CytK (2E-2D-CytK). We found that the greatest effect was obtained when the three rings of charges were most spaced [i.*e*., the (p(K)/p(Cl) was ~3.7 when the charges were placed between 1 and 2 nm apart from the charges at the *cis* and *trans* nanopore entry, **Figure** 1D]. The addition of a fourth charge increased the ion selectivity further, but only slightly (**Figure** 1E). Overall, the pore with the highest ion selectivity was K128D-Q145D-S151D-K155D-CytK (2E-4D-CytK, **Figure** 1F), which showed a (p(K)/p(Cl) of 4.04±0.07. At pH 3.8 the effect of the additional aspartate residues was minimal (**Figure** S1). An anion selective nanopore could also be made by replacing the negative charges for positive charges, resulting in the E112K-E139K-Q145K-S151K-CytK mutant (CytK-6K), with an anion selectivity of 0.207±0.008 (pH 7.5), which remained virtually the same at pH 3.8 (0.213±0.012).

To quantify the presence of EOF generated by the electroosmotic flow, we set up non equilibrium MD simulations of the WT pore and of the three mutants 2E-2D-CytK, K128D-K155D-Q145D-CytK (2E-3D-CytK) and 2E-4D-CytK (the system set-up is sketched in **Figure** S2). We found that at negative voltages, the EOF for the WT pore is directed from *trans* to *cis*, while for all other mutants is from *trans* to *cis* (**Figure** 1G). This change in the EOF direction is coherent with ion charge maps at equilibrium (*i.e*., ΔV=0, **Figure** 1H and **Figure** S3). In the WT pore, anions accumulate at the two extremities of the barrel in correspondence to K155 and K128 (**Figure** 1B). When these positively charged amino acids are exchanged into negative charges, the pore becomes cation selective. Cation accumulation is larger in pores bearing additional negative charges in the barrel region (**Figure** 1G). The dependence of the current and EOF from the voltage are roughly linear (**Figure** S4), confirming that the dominant mechanism for EOF is the presence of fixed charges at the wall surface, as shown by simulations of similar pores like αHL^15–17^ and not an induced charge effect as in ref^18^.

### Translocation of model substrates across CytK nanopores

The linearised translocation of a protein across a nanopore requires un-structuring the polypeptide and overcoming steric, entropic and electrostatic energy barriers. Unfolded polypeptide translocation was initially tested with S1, a 123 amino acid polypeptide designed to be unfolded and to carry large stretches of positive charges (at pH 7.5, S1 net charge is +28, or +23 charges every 100 amino acids also expressed as +23_100_, **Figure** 2A). The addition of S1 to the *cis* side of WT-CytK induced current blockades of two types (**Figure** 1B, S5), named type I and type II. Blockades were characterised by the duration of the blockade (dwell time) and the amount of excluded current I_ex%_ defined as I_ex%_ = (I_o_-I_b_)/I_o_ x 100, where I_b_ the amount of current during the polypeptide blockade and I_o_ is the open pore current. type I and type II blockade showed a complex voltage dependency. Likely, the two type of events reflects the translocation from the N- or C- terminus. The voltage dependency of the blockades indicates that the electrophoretic force drives S1 translocation across a non-selective nanopore (WT-CytK) above a threshold potential of ~ −160 mV and −140 mV for type I and type II events, respectively (see ref^20^ and references therein for additional discussion).

**Figure 2.**
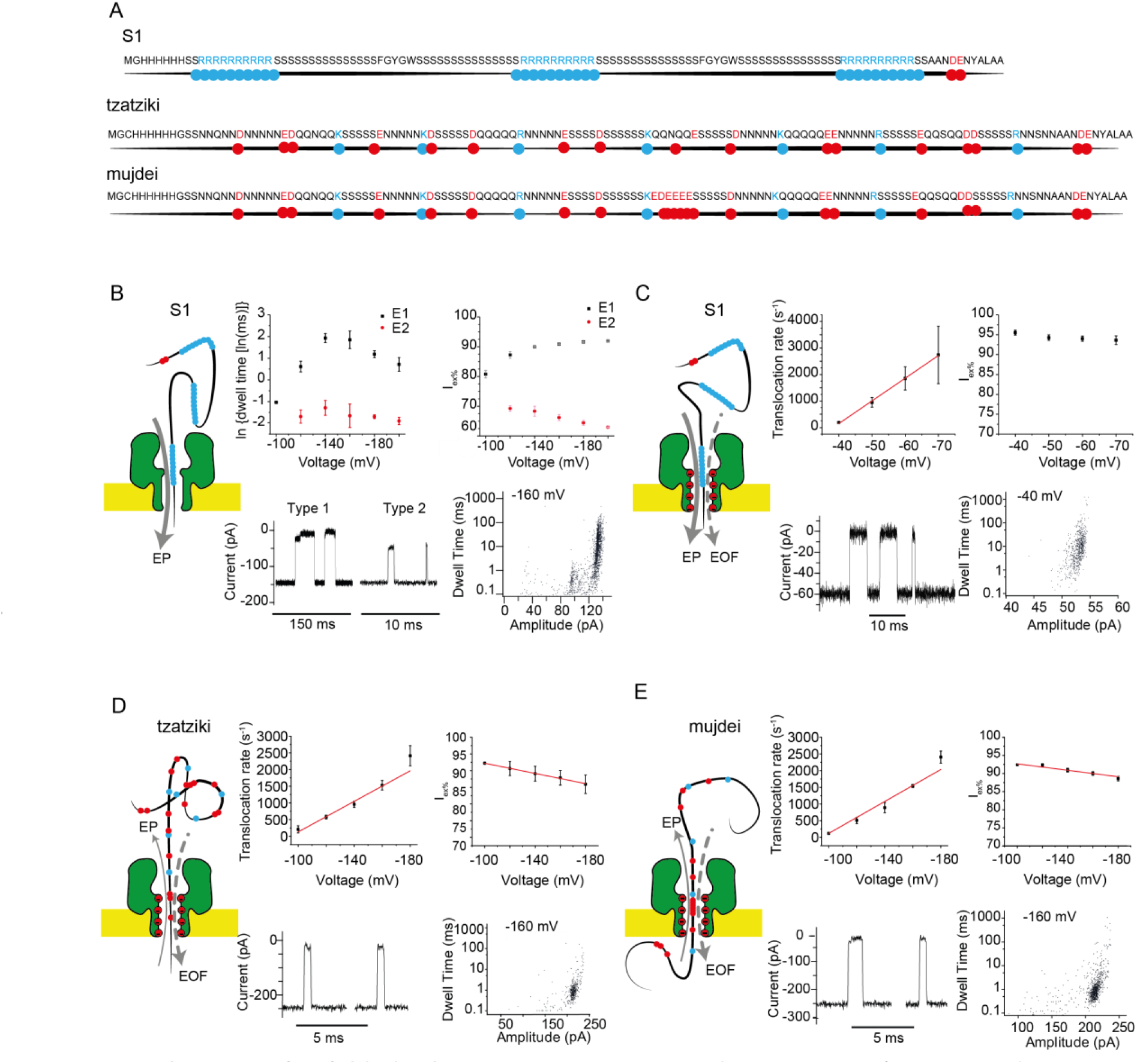
Translocation of unfolded substrates using engineered nanopores. A) Amino acid sequence and corresponding schematic representation of the three designed polypeptide substrates. Red dots indicate negatively charged amino acids; blue dots positively charged amino acids. B) S1 translocation through WT-CytK nanopores. The four panels from left to right and top to bottom indicate: the voltage dependency of the blockade duration, the voltage dependency of the excluded current (I_ex%_), representative traces at −160 mV, and the dwell time versus current amplitude at −160 mV. C-E) Translocation of the three substrates as indicated by the schematics on the left of the panel through 2E-4D-CytK nanopores. From left to right and top to bottom the panel shows the voltage dependency of translocation rate, the voltage dependency of the excluded current (I_ex%_), representative traces at the indicated bias, and the corresponding dwell time versus current amplitude at the indicated bias. Experiments were performed at pH 7.5 and under 1 M KCl. Traces were collected at 50 KHz sampling rates and filtered at 10 KHz using a Bessel filter.

Using the nanopores with enhanced electroosmotic flow the threshold bias to obtain translocation was strongly reduced (*e.g*., to ~-30 mV for 2E-2D-CytK), indicating that the EOF can further drive the translocation of the polypeptide. Interestingly, at low bias the nanopores with high charge density (*e.g*. 2E-4D-CytK) showed events lasting up to a few seconds (**Figure** 2C, **Figure** S6), which most likely resulted from the binding of stretches of positive charges in S1 to the negative charges within the nanopore. In general, however, above the threshold potential, the dwell time was strongly reduced by the additional EOF (**Figure** S7-S12). The distinction between type I and type II became less obvious (**Figure** S7) or inexistent (**Figure** S8-S11).

S1 is a highly positively charged model polypeptide, with an amino acid composition and distribution not found in native proteins. In order to test an unstructured native-like substrate and establish whether the engineered EOF strength can be used to translocate unfolded polypeptides against an EPF, which is a likely requirement if proteins are to be sequenced with nanopores, we prepared a new model polypeptide 140 amino acid long named tzatziki, illustrated in **Figure** 2A. Tzatziki was designed to be unfolded and to carry a relatively large negative charge density of −7.0_100_.

When Tzatziki was added to the *cis* side, blockades were not observed with the WT-CytK nanopores, indicating that the EPF is too weak to induce polypeptide translocation at up to +200 mV. In the case of 2E-2D-CytK, capture events were observed above −180 mV (substrate added to the *cis* side), but substrate translocation could not be proven by voltage dependencies (**Figure** S12). Tzatziki translocation events were observed in mutant pores with at least three rings of negative residues (cation selectivity >3.5, **Figure** S13-S15) at a bias above −120 mV (**Figure** 2D). The dwell time of tzatziki decreased with the EOF of the nanopore (**Figure** S16). Together these results indicate that the EOF drives the translocation of the polypeptide.

An EOF-driven translocation is also observed by MD simulations. Using the first 55 residues corresponding to the N-terminal fragment of Tzatziki (−4e over 55aa or −7.2_100_) threaded through 2E-4D-CytK, we found that at negative voltages the total force was directed from *cis* to *trans* in the direction of the EOF and opposite to the EPF which is directed in the opposite direction. The EOF is also stronger than the EOF also when using a peptide with a larger charge density (−9.1_100_, **Figure** S17). The differences in the total force acting on the peptide among the three replicas are related to the different conformations that the unfolded peptide acquires in the pore. This variability is expected since the hydrodynamic drag on an object strongly depends on its shape, even at similar average flow rate. Despite this variability, in all replicas the direction of the average force for negative voltages is from *cis* to trans, suggesting that the translocation of the peptide through the pore is unidirectional.

Proteins found in nature might contain stretches with a highly focused charge density. Since the EP drops very steeply within the nanopore and in particular near the constriction, we investigated whether a substrate with several consecutive negative charges could be translocated across 2E-4D-CytK nanopores. We designed mujdei, identical to tzatziki with the exception of a stretch of five negatively charged residues (EDEEE) in the middle region of the polypeptide (**Figure** 2A, S18). The overall charge density of this substrate increased to −9.9_100_. The 2E-4D-CytK nanopore could capture and translocate mujdei showing very similar dwell times compared to tzatziki, suggesting that the electroosmotic flow is considerably larger than the electrophoretic force. This work shows, therefore, that a relatively highly negatively charged polypeptide can be translocated against the EP.

In order to test whether an the EOF generated by anions could also have the same effect, we tested 6K-CytK nanopore a CytK-based nanopore where all the aspartate residues in the barrel of 2E-4D-CytK were substituted to lysine residues. Surprisingly, despite 6K-CytK nanopore as a large anion selectivity (p(Cl)/p(K) of 4.83±0.19 at pH 7.5), comparable in cation selectivity of 2E-4D-CytK, all the model substrates tested could enter but could not be translocated, nor easily released across the nanopore (**Figure** S19, note that a cis-to-trans EOF is generated at positive bias). A similar effect was observed with an engineered aerolysin nanopore (**Figure** S20), suggesting that this might be a generic effect for β-barrel nanopores.

### Stretching the polypeptides inside the nanopore

In nanopore protein sequencing it is important that the polymer is linearised during translocation and that there is enough current associated with the polypeptide blockade to identify individual amino acids. The I_ex%_ is typically dominated by the excluded volume of the polymer inside the nanopore. Therefore, if the polypeptide translocates as a linear polypeptide, the I_ex%_ is expected to be low, while if the polypeptide is folded inside the nanopore the I_ex%_ is expected to be high. However, it is important to note that the I_ex%_ might also be influenced by the charges of the analyte and the nanopore, which might create additional energy barriers for the translocation of ions from solution. For example, in a nanopore with a highly positively charged lumen the translocation of a polymer with high negative charge density such as DNA excludes the current almost completely, most likely because the transport of anions is blocked by the charge in the DNA and that of cations by the charge of the nanopore. Here, for most substrates, we observed a reduced I_ex%_ with the increasing of the applied voltage, suggesting that the polypeptide is stretched as the electroosmotic pulling force is increased. A notable exception was the translocation of S1 through 2E-2D-CytK nanopores (supporting information, **Figure** S6). Possibly, this effect is due to the charge distribution of S1 which might specifically interact with the fixed charges in the nanopore.

### Non-enzymatic translocation of native substrates across the 2E-4D mutant

S1, tzatziki and mujdei were designed to contain only disorder-promoting hydrophilic amino acids to minimise the folding of the polymer. In order to test the non-enzymatic translocation of native proteins, we chose a maltose-binding-protein malE219a (a variant containing the G220D and E221P destabilising mutations and a total of 412 amino acids, charge density of −2.3_100_ at pH 7.5, **Figure** 3A), and a glucose binding protein H152A-GBP (bearing a destabilising mutation H152A, 341 amino acids, charge density −1.1_100_ at pH 7.5, **Figure** 3A). As previously reported, malE219 fully unfolds in 0.7 M GuHCl^22^ while H152C-GBP is unfolded in 1 M GuHCl and 2 M urea, respectively.^23^

**Figure 3.**
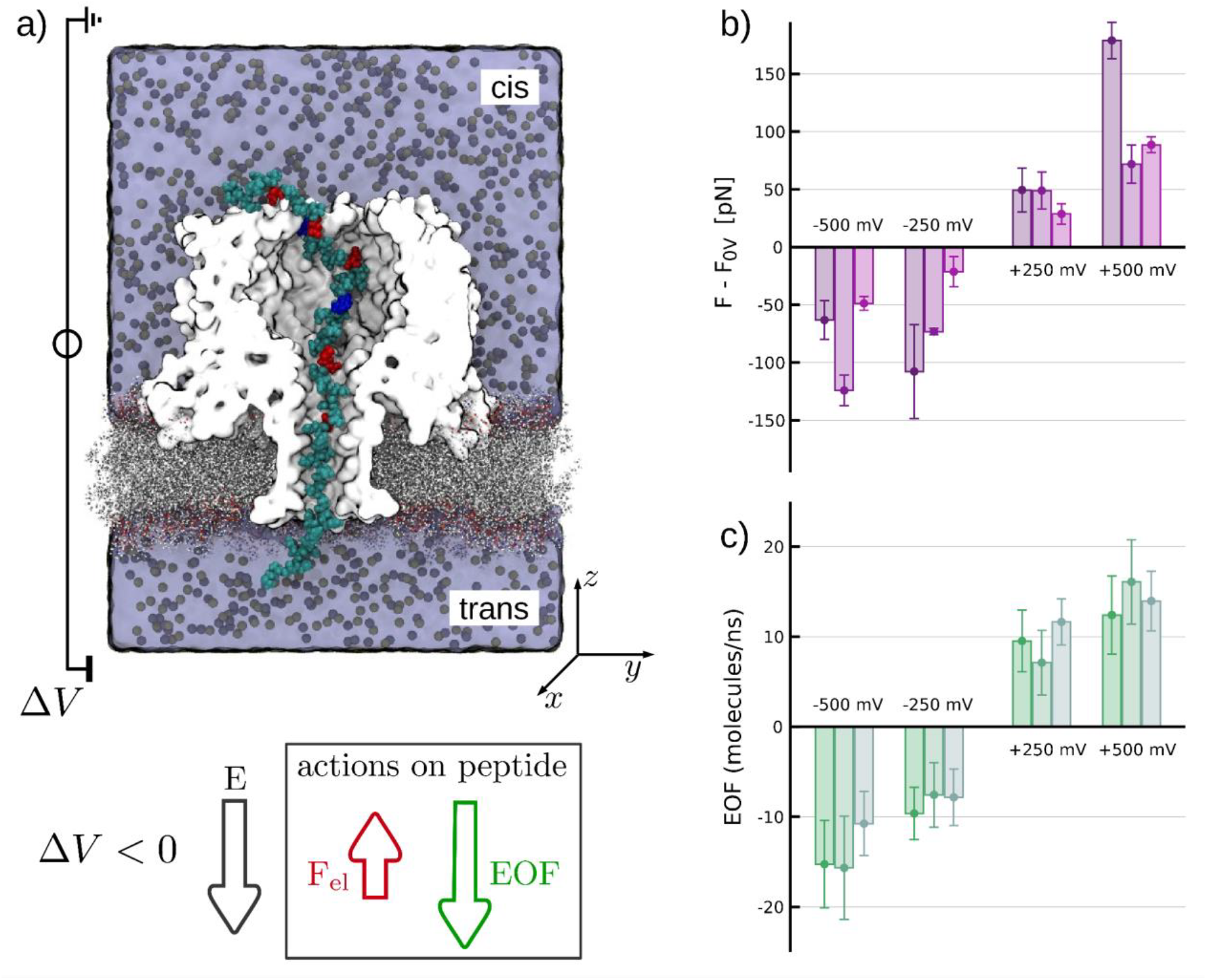
Forces on a Tzatziki fragment. **a)** Cut-through of one frame of the MD simulation showing the forces acting on the first 55 amino acids of Tzatziki, which was preliminary pulled into the nanopore with the N-term points toward the *trans* side. Water and ions inside the pore are not represented. The z-coordinate of the center of mass of the part of peptide spanning the beta barrel is constrained to its initial position with a harmonic spring, and a voltage is applied to the system. **b)** Net force on the peptide. The net force is shown as the difference in the average force along the axis in the presence (F) or absence (F_0V_) of an applied voltage. **c)** The EOF for the same simulations. Taken together, the resulting force is always directed as the EOF, indicating that the competition between the hydrodynamic drag on the peptide due to EOF and the electrophoretic force (F_el_) is dominated by the EOF. For negative ΔV, depicted in the lower left corner, this would result in a cis-to-trans movement. Confidence intervals for each run are estimated via block average. VMD software^21^ was used for panel a. Error bars represent the standard error of independent replicas.

The presence of urea (a neutral denaturant) or GuHCl (a charged denaturant) up to ~ 3M did not have a noticeable effect on the stability of the nanopore. Urea, however, decreased the open pore current of 2E-4D-CytK while retaining the current asymmetry under opposite bias (**Figure** S21A,C), suggesting that it did not change the ionic properties of the nanopore. By contrast, in the presence of Gu.HCl the current symmetry was lost and even reverted at higher concentrations (**Figure** S21B,D), suggesting that, as previously shown by MD simulations, the Gu.H^+^ binds to the negative charges within the lumen of the nanopore,^1^ thus reducing the ion selectivity of the 2E-4D-CytK nanopore.

When adding (pre-unfolded in 1.5 M GuHCl) malE219a substrate to the *cis* side of 2E-4D-CytK in the presence of 2 M urea, we observed translocation events, which were significantly different from the translocation of the model substrates (**Figure** 3A, S22). The threshold translocation of malE219a was ~-60 mV compared to ~-100 mV for tzatziki, possibly reflecting the lower negative charge density of the protein (−2.3_100_ vs −7.0_100_). The dwell times, on the other hand, was about two-fold longer compared to the dwell time of the model substrates adjusted for their length, possibly reflecting a stronger interaction between the hydrophobic amino acids and the lumen of the nanopore. Importantly, during the blockade, a substantial larger residual current was observed during the translocation of the protein (I_ex%_ 70.5±0.57% at −120 mV (**Figure** 3A) vs 88.79%±2.03 (tzatziki) or 90.05±0.50 (mujdei) at −160 mV, **Figure** 2D-E), suggesting a more stretched polypeptide. Furthermore, the current signature showed rather distinctive patterns, which could be related to the sequence or, more likely, to the residual structure of the proteins during translocation.

Pre-unfolded H152A-GBP (added to the *cis* side) also translocated through 2E-4D-CytK nanopores in the presence of urea (2.44 M), although the threshold for translocation was higher than for malE219a (−80 mV, **Figure** 4B, S23) despite having a lower charge density (−1.1_100_ vs −2.3_100_). Interestingly, the I_ex%_ showed two consecutive levels: level 1 defined by an I_ex%_ of 70.62±1.08% (at −120 mV), which is similar to the I_ex%_ measured for malE219a, followed by Level 2 with an I_ex%_ of 89.11±3.50% (at −120 mV, **Figure** 4B), which is similar to the I_ex%_ of model substrates. This behaviour may be explained by the two-step unfolding of H152A-GBP, as previously observed for thioredoxin,^2^ or by the translocation of a partially folded structure.

**Figure 4.**
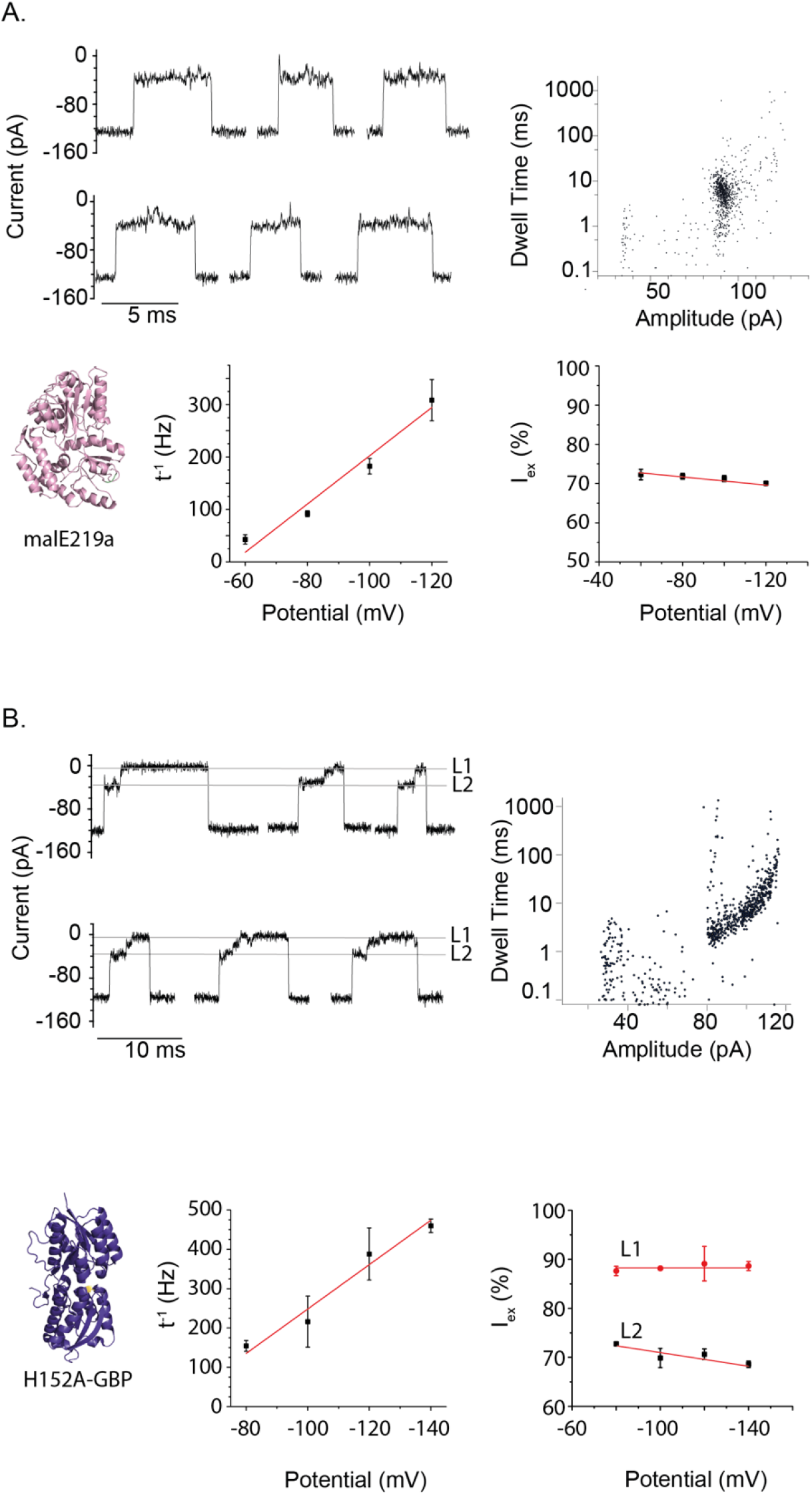
Translocation of unfolded proteins across 2E-4D-CytK nanopores. A) Translocation of MalE219a across 2E-4D-CytK in 2 M urea. Top, representative traces (left) and dwell time versus amplitude (right) under −100 mV. Bottom, cartoon representation of MalE219a (left), voltage dependency of the translocation speed (centre), and voltage dependency of the excluded current (right). (B) Translocation of H152A-GBP across 2E-4D-CytK in 2.4 M urea. Top, representative traces of unfolded H152A-GBP translocations showing Level 1 (L1) and level 2 (L2, left), and dwell time versus amplitude (right) under −100 mV. Bottom, cartoon representation of H152A-GBP (left), voltage dependency of the translocation speed (centre), and voltage dependency of the excluded current (right). Experiments were performed at pH 7.5 and under 1 M KCl. Traces were collected at 50 KHz sampling rates and filtered at 10 KHz using a Bessel filter.

When GuHCl (1 M or 1.8 M) was used with 2E-4D-CytK, we observed translocation events at negative bias, but with a reduced frequency and the threshold potential shifted towards higher values. Hence, the EOF induced by the nanopore is reduced, most likely due to the binding of GuH^+^ ions to the nanopore,^1^ but it still allows polypeptide translocation against the EP. We also found that in 2 M urea the I_ex%_ in urea was substantially lower than in 1.8 M GuHCl (70.5±0.57% vs 88.75±0.31% at −120 mV) and the dwell time longer (3.28±0.45 ms vs 1.90±1.02 ms, **Figure** S25). A possible explanation is that the polypeptides in the presence of urea are more stretched (higher I_ex%_) due to the stronger EOF, or that the binding of GuH^+^ ions to the nanopore reduce the volume for peptides translocation.

Notably, previous work showed that unfolded proteins might be captured by β-barrel nanopores such as α-hemolysin (homologous to CytK) or aerolysin, in the presence of denaturants.^11–13,24,25^ Some work reported the translocation of unmodified proteins,^24,25^ while other attached either a ssDNA^11,12^ or a poly-10 aspartate tag^13^ at the N- or C- termini to drive the EPF-driven transport across nanopores. We tested malE219a and malE219a-D10ssrA (malE219a expanded with a D10ssrA tag at the C-terminus) in the presence of 1.5 M GuHCl and using WT-CytK. We found that while malE219a could not be transported at any tested bias (up to +200 mV); malE219a-D10ssrA translocated across the nanopore at positive applied potentials (**Figure** S24). We conclude, therefore, that the EOF induced by Gu.HCl alone cannot drive polypeptide translocation, but an additional EPF force on the D10 tag must be applied, at least to induce the initial threading of the protein. By contrast, our work shows that by introducing fixed charges on the nanopore’s wall, an EOF can be engineered to translocate native proteins and to overcome opposing EP during translocation.

## Conclusion

The sequencing of protein using nanopores requires that proteins are unfolded, using enzymes or denaturants, and translocated across the nanopore, possibly amino acid by amino acid. Furthermore, the polypeptide should be linearised during translocation so that individual amino acids can be addressed. Contrary to nucleic acids, polypeptides do not have a uniform charge distribution, and their transport and linearisation cannot be induced by the electric field inside the nanopore. In this work, we found that an EOF can be engineered to drive the translocation of polypeptides and proteins against the electrophoretic force, which is a fundamental requirement for both enzymatic or non-enzymatic protein sequencing with nanopores.

Using CytK, we found that at least three rings of negatively charged residues, preferably equally spaced, must be added along the 5 nm lumen of the nanopore to induce an EOF that allows the transport of unfolded polypeptides. Surprisingly, the introduction of positively charged did not induce the same effect. Possibly either the size of introduced amino acid or the nature of the counterion are important factors to allow protein translocation.

This approach can be used in the presence of denaturants to induce the breaking of the folded structure of proteins and the concomitant electroosmotic-driven transport of the unmodified protein across the nanopore. Remarkably, using 2E-4D-CytK the nanopore with the strongest EOF, we also showed that proteins with a large negative charge density (9.9_100_) can be transported through the nanopore against the EPF exerted on the polypeptides during translocations.

Non-enzymatic protein transport across nanopores might offer fast and high-throughput means to characterise single molecules. Here, we show that urea might be used to unfold proteins, while the electroosmotic flow generated by the surface charge of the nanopore is used to induce the translocation of proteins. Urea is preferred to the use of GuHCl, as it allows the transport of larger ionic currents. In nanopore protein sequencing, enzymes will probably be used to control the unfolding and precise speed of polypeptide translocation while individual amino acids are identified as they pass a recognition site inside the nanopore. The engineered EOF will allow the capture and translocation of all polypeptides without requiring the use of electrophoretic tags. Finally, the EOF will also stretch the polypeptide during translocation, which will be important to allow the recognition of individual amino acids as they pass the recognition site.

## Ackomedgment

The authors acknowledge supercomputer time provided through s1178 Production Grants by CSCS, NOW-VICI (grant 192.068) and NIH (grant 1086554). M.C and B.M.D.R thanks Giovanni Di Muccio for useful discussion.

