## Supplemental information for "An engineered electroosmotic flow transports unravelled proteins across nanopores"

### Materials and Methods

#### Chemicals and reagents

The chemicals and suppliers used is listed as follow: Amplicillin sodium salt was purchased from Fisher Bio Reagents; chloramphenicol (≥98.0) from Sigma Life Science); urea (≥99.5%), guanidinium chloride (≥99.5%, biochemistry), isopropylthio-β-galactoside (≥99.0%, dioxin-free, animal-free), LB medium, 2xYT medium, NaCl (≥99.5%), HEPES (PUFFERAN® CELLPURE® (≥99.5%), imidazole (≥99%), KCl (≥99.5%), Tris(2-carboxyethyl)phosphine hydrochloride (≥98.0%), Dodecyl-β-D-maltosid (≥99%) from Roth; citric acid (≥99.6%, anhydrous and n-hexadecane (99% from Acros Organics; BIS-TRIS propane (≥99.0%) from Sigma Life Sciences; protease inhibitors (Pierce™ Protease inhibitor Mini tablets, EDTA-free); GeneJET gel extraction kit, GeneJET PCR purification kit, GeneJET Plasmid Miniprep kit were purchased from (Thermo Scientific); Ni-NTA agarose from Qiagen; Strep Tactin® Sepharose® and D-desthiobiotin from IBA Lifesciences; mPEG-mal 5k from Laysan Bio, Inc.; DPhPC from Avanti polar lipids, n-pentane from Sigma-Aldrich; Quick Start Bradford 1x Dye Reagent (Bio-Rad); DNA primers and gBlock™ from IDT. BL21(DE3) strain harbouring the pET-PfuX7 plasmid was kindly provided by prof. dr. Oscar Kuipers

#### Methods

*Cloning and production of the CytK mutants*

Plasmids containing the mutant CytK nanopores were constructed by means of USER cloning[1,2] using uracil-containing primers. In short, the full gene was split into upstream and downstream fragments at the mutation site, and a homology region of 8-13 bps was defined around this site. The empty Pt7-SC1 backbone was also amplified with uracil-containing primers. The upstream (F1) and downstream (F2) fragments would be rejoined together with the empty vector (V) in the USER reaction. PfuX7 DNA polymerase was produced as previously described[3] and used to generate the two gene fragments and the linearised pT7-sc1 (AmpR). The PCRs was performed as in ref[3], with the exception of the extension time, which was lowered from 1 min/kb to 30 s/kb. The PCR products were either gel extracted (if by-products were present), or directly cleaned up from the PCR mix. The gene fragments and the linearised vector were mixed in a molar ratio F1:F2:V of 3:3:1 and the USER reaction was performed as in ref[2]: 25 min at 37 °C, followed by a 60 °C incubation step of 10 min. Lastly, the mixture was cooled down to RT (22 or 20 °C) for 15 min and subsequently stored on ice/ at 4 °C until the transformation step. The circularised plasmids were transformed into chemically competent *E. cloni* cells using the heat shock procedure and the cells were selected on a LB-agar plate supplemented with 100 µg/mL ampicillin. Plasmids from individual colonies were isolated and the introduction of the mutations was confirmed by Sanger sequencing (Macrogen).

The plasmids encoding for the CytK mutants were electroporated into BL21(DE3) electrocompetent cells using a Bio Rad Micro Pulser (bacterial setting), and the cells were selected on plates containing 100 µg/mL ampicillin. Next day, several transformants were resuspended in LB medium supplemented with 100 µg/mL ampicillin and added to 200 mL LB medium containing 100 µg/mL ampicillin such that the starting optical density at 600 nm ($OD_{600}$) was 0.05-0.1. Cells were grown at 37 °C, 180 RPM until an $OD_{600}$ of 0.6-0.8, when the culture was chilled on ice for 5-10 min, followed protein expression being induced with 0.5 mM IPTG. After an incubation of 19-21 h at 25 °C, 180 RPM, the cells were harvested (7500 rpm, 5 min). Following a 1 h incubation at -80 °C, the cell pellets (100 mL culture) were resuspended in 20-25 mL ice-cold lysis buffer (50 mM HEPES, 150 mM NaCl, 10 mM imidazole, pH 7.4 + 0.02% DDM) supplemented with ¼ tablet cocktail protease inhibitors per 100 mL cell culture and subsequent steps were performed at 4 °C degrees unless stated otherwise. The cell suspension was sonicated (Branson sonifier 450) at 25% duty cycle, 2.5 output control for 2-3 min and the cellular debris was removed (8000 RPM, 20 min). The supernatant was incubated (with shaking) for 20-30 min with 200 µL $Ni^{2+}$-NTA slurry (50 % suspension), pre-equilibrated and prewashed with 1 mL lysis buffer. The beads were briefly pelleted (3000 RPM, 1 min) and transferred to the column (1.2 mL bed volume bio-spin chromatography, BioRad) while allowing the flow through pass, at RT The column was washed in steps with 10 mL wash buffer (50 mM HEPES, 150 mM NaCl, 30 mM imidazole, pH 7.4 + 0.02 % DDM). The protein was eluted with 200 µL elution buffer (50 mM HEPES, 150 mM NaCl, 250 mM imidazole, pH 7.4 + 0.02% DDM) in three elution fractions. The presence of the SDS-stable CytK mutant oligomers was confirmed by SDS-PAGE, omitting the heating step in the sample preparation.

*Preparation of the protein substrates*

The DNA sequence of the S1 substrate was previously prepared in our lab.[4] The plasmid bearing tzatziki was prepared from a gBlock™ introduced in the linearised pT7-sc1 using USER cloning as previously described. The DNA for mujdei was generated by using tzatziki as template. Fusion PCR was used to obtain the full mujdei insert, which would be introduced into pT7 vector also by USER cloning. The malE219a, malE219 and malE219aD10ssrA substrates were prepared in a similar fashion. All substrates' sequences were confirmed sequencing (Macrogen). The substrates were expressed in the SG1146a strain in order to limit protein degradation. The plasmids containing the S1 and malE219a DNA were transformed into chemically competent SG1146a cells and transformants were selected on ampicillin-containing plates. Next day, several colonies were resuspended in LB medium with ampicillin and added to the culture medium (LB supplemented with 100 µg/mL Amp and 25 µg/mL Chloramphenicol) such that the starting OD600 was 0.05-0.1. Cells were grown until OD600 0.6-0.8, when the culture was chilled (5-10 min on ice) followed by induction with 0.5 mM IPTG at 25 C, 180 rpm for 18-22 h. The cells were harvested (7500 rpm, 5 min) and stored at -80 C for 1 h, followed by resuspension in 20 mL lysis buffer (50 mM HEPES, 150 mM NaCl, 10 mM imidazole, 6 M GuHCl, pH 7.4) at RT and all the subsequent steps were performed at RT. The cell suspension was sonicated (25% duty cycle, 2.5 output control for 2-3 min). The cellular debris was removed (8000 RPM, 20 min) and the resulting supernatant was incubated with 200 µL $Ni^{2+}$-NTA slurry per 100 mL culture with shaking for 30 min. The resin was transferred to the column (2 mL, biorad) and washed with 10 mL wash buffer (50 mM HEPES, 150 mM NaCl, 30 mM imidazole, 1.5 M GuHCl, pH 7.4. Lastly, the protein was eluted four times in 100 µL elution buffer (50 mM HEPES, 150 mM NaCl, 250 mM imidazole, 1.5 M GuHCl, pH 7.4). The presence of the protein was confirmed by SDS-PAGE, followed by aliquoting of the elution fractions and storage at -20 C. GBP H152A protein was kindly provided by Nicole Galenkamp. The protein concentration was determined by Bradford assay.

The plasmids containing the tzatziki and mujdei plasmids were transformed into electrocompetent SG1146a cells, which were plated on Amp plates. Several transformants were resuspended in 2YT medium and diluted into 2YT medium supplemented with 100 µg/mL Amp and 20 µg/mL Cam to a final OD600 of 0.05-0.1. When OD600 reached 0.6, protein expression was induced with 0.5 mM IPTG at 25 C, 180 rpm for 18-20 h. The proteins were purified as described for S1 and malE219, although under native conditions (thus omitting the GuHCl in the buffers). The presence of the protein was confirmed by SDS-PAGE and using a PEG-maleimide reaction as an additional check, while Bradford assay was insensitive to the presence of these substrates (due to their amino acid composition).

*Electrophysiology measurements*

Recordings in planar lipid bilayers were performed using a chamber consisting of two compartments, delimitated by a 25 µm thick Teflon membrane which an aperture of approximately 100 µm as described earlier.[5] In short, a droplet (half the quantity contained in a 10 µL glass capillary) consisting of n-hexadecane dissolved in n-pentane (6.25%) was applied on the Teflon membrane, followed by the addition of 500 µL buffer and two droplets of DPhPC lipids in n-pentane (5 mg/mL) in each compartment. Ag/AgCl electrodes were connected to each chamber via agarose bridges (2.5% agarose, 3M KCl solution), grounding the *cis* compartment. Measurements were performed using an Axon™ Digidata® 1550B digitizer and an Axopatch 200B amplifier (Molecular Devices) and recorded with the Clampex 11.1 software.

*Ion selectivity*

Buffers used: Buffer A - 2 M KCl, 15 mM HEPES, pH 7.5, Buffer B - 0.5 M KCl, 15 mM HEPES, pH 7.5, Buffer C - 0 M KCl, 15 mM HEPES, pH 7.5, Buffer D - 2 M KCl, 50 mM citric acid, Bis-tris propane (BTP), pH 3.8, Buffer E - 0.5 M KCl, 50 mM citric acid, BTP, pH 3.8, Buffer F - 0 M KCl, 50 mM citric acid, BTP, pH 3.8

The ion selectivity of the CytK mutants was determined in buffers of either pH 7.5 or 3.8. Firstly, both the *cis* and *trans* compartments were filled with 2 M buffer (Buffer A or Buffer D) and a single nanopore was isolated, followed by the pipet-offset adjustment of the current at 0 mV bias to 0 pA. The I/V curve was determined between -140 mV and +140 mV, in steps of 20 mV, using a 10 kHz sampling rate coupled with a 2 kHz Bessel filter. Next, the concentration of KCl in the *trans* compartment was lowered to approximately 0.5 M by flushing with 0 M buffer (Buffer C/ Buffer F) and repeated flushing with 0.5 M buffer (Buffer B/ Buffer E) ensured that the final concentration of KCl in the trans compartment was correctly fine-tuned to 0.5 M. Similarly, the I/V curve of the nanopore was measured. The reversal potential was determined from the second I/V curve from the linear function fitting the data points between -20 and +20 mV. The ion selectivity, expressed as the fraction $p_{K+}/p_{Cl-}$ was calculated using the formula below and each ion selectivity was established from triplicate experiments.

$$\frac{p_{K+}}{p_{Cl-}} = \frac{[a_{Cl^-}]_{trans} - [a_{Cl^-}]_{cis} \times e^{V_r F/RT}}{[a_{K^+}]_{trans} \times e^{V_r F/RT} - [a_{K^+}]_{cis}}$$

where [a] is the activity of the $K^+$ or $Cl^-$ in the *cis* or trans compartment, $V_r$ is the reversal potential, which is obtained from the experiments, F corresponds to the Faraday constant (96 485 C/mol), R the gas constant (8.3145 J $mol^{-1}$ $K^{-1}$) and T the temperature (298 K).

*Translocation experiments*

Single pores of the CytK mutants were isolated and the pore orientation was determined from the I/V curve. Provided the pore vestibule was located in cis, the model substrates were added in the *cis* compartment (30 nM final concentration for S1 or 5-7 µL tzatziki or mujdei from the elution sample) and translocation was induced by applying a negative bias. These recordings were collected at 50 kHz sampling rate and a Bessel filter of 10 kHz, using a sweep protocol where the first 200-500 ms were used to unclog the pore by applying a positive bias, followed by approximately 2 s of recording at negative bias. In the case of native substrates, urea was introduced into the system, after determining the pore orientation and prior to substrate addition, by flushing both *cis* and *trans* compartments with 1 M KCl, 4 M urea, 15 mM HEPES, pH 7.5 buffer in 100 µL steps until the aimed concentration was reached (e.g. 2 M urea). After addition of urea, the substrate was added in the *cis* compartment in the following final concentrations: 0.3 uM malE219a, 0.4 uM H152A-GBP, 0.2 uM malE219aD10ssrA; translocation was followed like described for the other substrates. Each individual set of conditions was tested in triplicate.

*Data analysis*

In the case of the translocation experiments, the files containing the recorded sweeps were analysed using the Clampfit 11.1 software. Firstly, the open pore current (level 0, L0) and the corresponding noise, σ, was determined from the conventional histogram. Secondly, the detection limit, L1, was set at 10σ, (not that Clampfit sets it half-way detection algorithm at 5σ). Event detection with the set L0 and L1 was done on those approximately 2 s of recording at negative bias, with a dwell time cut-off of 0.08 ms. The resulting L1 data points were used to construct the log(dwell time) vs amplitude scatter plot, from which the amplitude boundaries of the cluster are defined. Next, using the amplitude boundaries, the logarithmic histogram of the dwell time and the conventional histogram of either the amplitude or the $I_{ex\%}$ are constructed. In both cases, the bin value was set such that the distribution within the histogram would resemble as much as possible a Gaussian shape. The values for the log(dwell time) and either the amplitude or the $I_{ex}$ were established by fitting a Gaussian function to the histogram, whose μ is either log(dwell time) or the amplitude/$I_{ex\%}$.

### Model of the Cytk pore

Cytk sequence (Uniprot ID Q937V0) devoid of the N-terminal signal sequence 1-31 was used to model a monomeric structure by feeding it to the homology model server SWISS-MODEL,[6] using the pore structure of alpha-hemolysin as a template (PDB ID 3ANZ[7]). As the model does not confidently assign initial residues, N-terminal modeling was performed as follows. Cytk monomer sequence was blasted to find homologues and structures from Leukotoxin LukEv, (PDB ID 7P8S,[8] and PDB id 3ROH[9]), along with the predicted Alphafold[10] structure (AF-Q937V0-F1), were used as templates to model the extra residues at the N-terminal. The missing residues MAQTT were added using the backbone coordinates of the beta hairpin region of 7P8S up to Val6, and prolonging the resulting beta strand. Sidechains of residues -1 to 6 were adjusted manually to minimize clashes, followed by a round of minimization with UCSF Chimera (Vers. 1.14,[11] 100 steps of steepest descent, 10 conjugate gradient, step size 0.02 Å). To fit the extra volume in the multimeric assembly, the loop Q95-S99 was adjusted via the Autosculpting module of PyMol [PyMOL]. The resulting structure was subject to another round of minimization with UCSF Chimera (same parameters). The resulting monomer was then mapped on the heptameric starting structure using the MatchMaker tool of UCSF Chimera. Single chain mutants were obtained using VMD[12] mutate function on corresponding residues prior to assembly in the heptameric channel.

### General Molecular Dynamics Simulation Methods

MD runs were performed using NAMD,[13] with CHARMM36 force field.[14] The water model used is TIP3P[15] and non-bonded fix corrections were applied for ions.[16] A time step of $\Delta t$ = 2.0 fs was used and long-range electrostatic interactions were evaluated via particle mesh Ewald method[17] with a 1 Å spaced grid. A cutoff of 12 Å with a switching distance of 14 Å was set for the short-range non-bonded interactions and periodic boundary conditions along the three directions were imposed. All covalent bonds with hydrogen were kept rigid, using SETTLE[18] for water molecules and SHAKE/RATTLE[19] for the rest of the system. A Langevin thermostat was used for all the simulations and a Nosé–Hoover Langevin piston pressure control was used for constant pressure simulations during the equilibration steps.[20]

*Pore and membrane system set-up and equilibration*

The modeled heptameric Cytk pore, aligned along the z axis, was inserted in a POPC bilayer membrane, parallel to the x-y plane, with a procedure already described,[21,22] and simulation box was completed with water and KCl at the final concentration of 1M plus extra ions to neutralize the system. Energy minimization and system equilibration was obtained with the protocol described in[23]. In brief, after minimization, a first equilibration step (P=1 atm, T=300 K) is performed to let the lipid tails melt and the electrolyte relax. External forces were applied to the water molecules to avoid their penetration into the membrane, while the backbone of the protein and the lipid heads were harmonically constrained to their initial positions. A second equilibration run was performed to compact the membrane reducing the spring constant on the protein backbone and removing the constraints on lipid heads. The last equilibration step is an NPT run without constrains. At the end of the equilibration procedure, the periodic box has approximately the following size: Lx = Ly = 137 Å, and Lz = 185 Å, for a total of around 360 000 atoms.

*Non-equilibrium MD simulations*

Production runs were performed applying a constant electric field whose component along the z coordinate is Ez. This is equivalent to a voltage drop of ΔV=EzLz, where Lz is the height of the periodic box.[24] Lipid head phosphorus and protein Cα atoms are harmonically constrained with spring constant kb = 10 kcal/(mol Å2). A thermostat (T = 300 K) is applied to the lipid and protein atoms. Ionic and electroosmotic currents were obtained with standard protocols reported in [10.1016/S0006-3495(01)75946-2, 10.1529/biophysj.104.058727, 10.1007/s10404-017-1928-1]. Block averages were used to estimate uncertainties. 3D ion concentration and charge density fields were calculated via VMD Volmap [10.1016/0263-7855(96)00018-5], on a cubic grid of side 1Å averaging over the stationary state of the non-equilibrium runs. The 3D Cartesian fields were mapped to a cylindrical coordinate system (r, z, θ) and averaged on θ to get density and velocity fields in the (r, z) plane.

*Peptides inside the pore*

Peptides were generated with Molefacture package of VMD[12] and pre-equilibrated in a suitable triperiodic box with water and ions for about half a ns. Final structures were then placed on the *cis* side of the equilibrated pore-membrane system, aligned with the pore axis, with peptide N-terminal about 2 nm above the vestibule opening. Water and ion molecules overlapping with the peptide were removed and the system was neutralized and further equilibrated (NPT flexible cell-constant area for 0.1 ns, P=1 atm, T=300 K) to stabilize Lz. Next, for the endecameric peptides, an NVT Steered MD with constant velocity (harmonic spring constant k=1 kcal/(mol Å2) ) was employed to bring the N-terminal of the peptide inside the beta barrel of CytK, reaching a position at ~0.5 nm from the trans side opening. The SMD was applied to the Ca of the N-terminal residue and velocity was set so that the peptide insertion is achieved in a 10 ns run. Typical velocities were around 1.2 nm/ns. Then the z-coordinate of the center of mass of the peptide was constrained with a harmonic spring (k=1 kcal/(mol Å2) and an additional 1 ns run was performed to further relax the system. The resulting configuration was used as initial condition for non-equilibrium runs with different applied voltages, where the peptide center of mass is constrained with the same spring. During the SMD and the applied voltage runs, the lipid heads of the membrane were harmonically constrained to their initial positions. The force acting on the peptide were obtained from the displacement of its center of mass. The protocol was repeated, starting from the peptide insertion step, for each replicate. The typical duration of production runs were between 40 and 70 ns, depending on the applied voltage, with the first 10 ns discarded in the analysis. A similar procedure was used for the 55aa peptide corresponding to the N-terminal portion of Tzatsiki; in this case only the center of mass of the residues inside the barrel is constrained in the production runs.

### Tables

*Tabel 1. Charge densities of investigated proteins*

| protein | charge at pH 7.5 | length (AAs) | charge density |
|---|---|---|---|
| malE219a | -9.4 | 412 | -2.28155 |
| GBP (Ec) H152A | -3.6 | 341 | -1.05572 |
| S1 | 28.3 | 123 | 23.00813 |
| tzatziki | -9.8 | 140 | -7 |
| mujdei | -13.8 | 140 | -9.85714 |

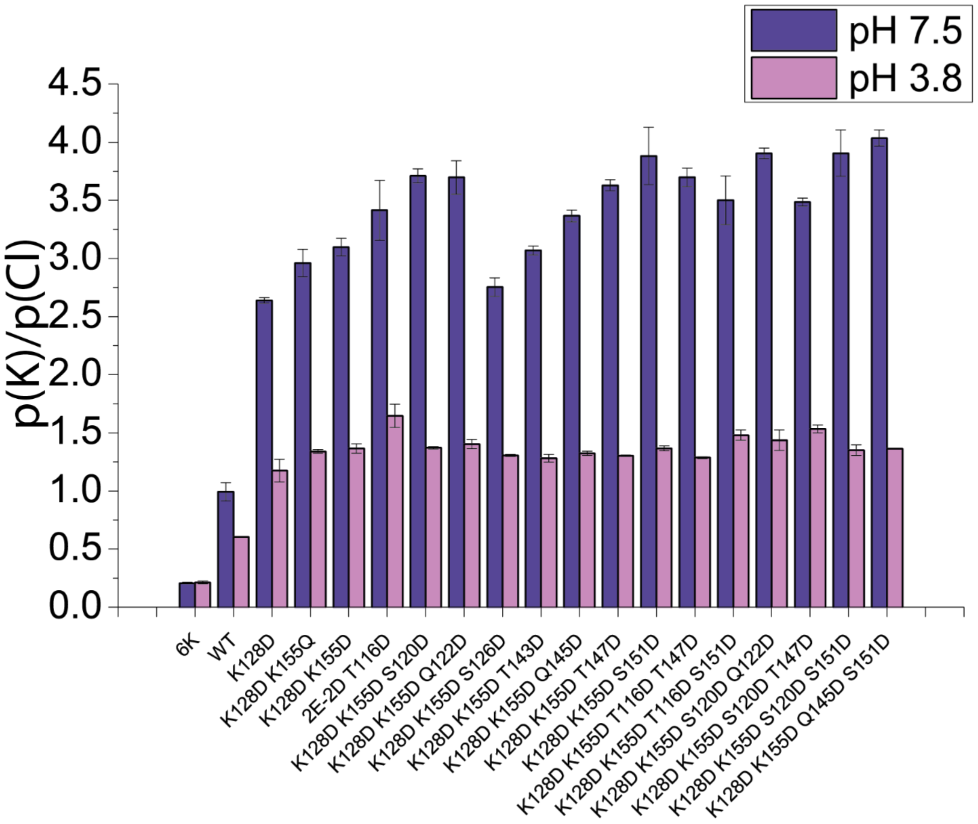

| | pH 7.5 | | | | | pH 3.8 | | | | |
|---|---|---|---|---|---|---|---|---|---|---|
| Mutant | R.P. avg | R.P. stdev | $P_K/P_{Cl}$ avg | $P_K/P_{Cl}$ stdev | | R.P. avg | R.P. stdev | $P_K/P_{Cl}$ avg | $P_K/P_{Cl}$ stdev | abbreviation CytK |
| WT | -0.14499 | 0.93088 | 0.991853 | 0.078977 | | -6.9109 | 0.090114 | 0.606714 | 0.004992 | WT |
| K128D | 12.90689 | 0.110296 | 2.637693 | 0.02426 | | 2.1993 | 1.201764 | 1.174445 | 0.097804 | 2E-1D |
| K128D K155Q | 14.24263 | 0.452406 | 2.959448 | 0.118611 | | 4.082796 | 0.168512 | 1.339853 | 0.016365 | 2E-1D-1Q |
| K128D K155D | 14.77041 | 0.276684 | 3.097926 | 0.075129 | | 4.333412 | 0.4065 | 1.364928 | 0.039817 | 2E-2D |
| K128D K155D T116D | 15.80865 | 0.829636 | 3.414994 | 0.2565 | | 6.874349 | 0.674812 | 1.645894 | 0.099883 | 2E-2D-T116D |
| K128D K155D S120D | 16.73486 | 0.17531 | 3.710497 | 0.061888 | | 4.402462 | 0.094033 | 1.37116 | 0.009331 | 2E-2D-S120D |
| K128D K155D T147D | 16.50404 | 0.137101 | 3.629647 | 0.046791 | | 3.700666 | 0.071011 | 1.303281 | 0.006666 | 2E-2D-T147D |
| K128D K155D S151D | 17.17087 | 0.673547 | 3.880897 | 0.247497 | | 4.346843 | 0.218138 | 1.365795 | 0.021517 | 2E-2D-S151D |
| K128D K155D S120D S151D | 17.2489 | 0.529757 | 3.906139 | 0.198708 | | 4.18201 | 0.379784 | 1.349959 | 0.045391 | 2E-2D-S120D-S151D |
| K128D K155D T116D T147D | 16.70064 | 0.180842 | 3.698435 | 0.077535 | | 3.520646 | 0.075161 | 1.286474 | 0.008517 | 2E-2D-T116D-T147D |
| K128D K155D T116D S151D | 16.10496 | 0.535072 | 3.501811 | 0.209061 | | 5.44246 | 0.335793 | 1.479538 | 0.044699 | 2E-2D-T116D-S151D |
| K128D K155D S120D T147D | 16.07314 | 0.08806 | 3.485588 | 0.035042 | | 5.92812 | 0.24134 | 1.53296 | 0.033551 | 2E-2D-S120D-T147D |
| K128D K155D S126D | 13.41838 | 0.276886 | 2.75487 | 0.079607 | | 3.729563 | 0.070704 | 1.306001 | 0.008162 | 2E-2D-S126D |
| K128D K155D T143D | 14.66842 | 0.115902 | 3.069036 | 0.038374 | | 3.465806 | 0.2951 | 1.281696 | 0.033533 | 2E-2D-T143D |
| K128D K155D Q122D | 16.69164 | 0.331709 | 3.697167 | 0.144758 | | 4.722214 | 0.275372 | 1.403646 | 0.03974 | 2E-2D-Q122D |
| K128D K155D Q145D | 15.69622 | 0.133073 | 3.36655 | 0.050095 | | 3.905657 | 0.158716 | 1.32279 | 0.018525 | 2E-2D-Q145D |
| K128D K155D S120D Q122D | 17.26572 | 0.094966 | 3.905306 | 0.044557 | | 5.022353 | 0.692365 | 1.436375 | 0.088142 | 2E-2D-S120D-Q122D |
| K128D K155D Q145D S151D | 17.60126 | 0.139616 | 4.036893 | 0.068788 | | 4.313304 | 0.014851 | 1.362292 | 0.001796 | 2E-4D |
| E112K E139K Q145K S151K | -19.3082 | 0.295947 | 0.20764 | 0.008135 | | -19.0541 | 0.430281 | 0.213452 | 0.012095 | 6K |

**Figure S1. CytK ion selectivity.** Top: Graph of the cation selectivity measured for the CytK mutants at pH 3.8 and 7.5. With the exception of the WT Bottom: The table containing the values determined for the p(K)/p(Cl) of the mutants. All measurements were done in triplicate.

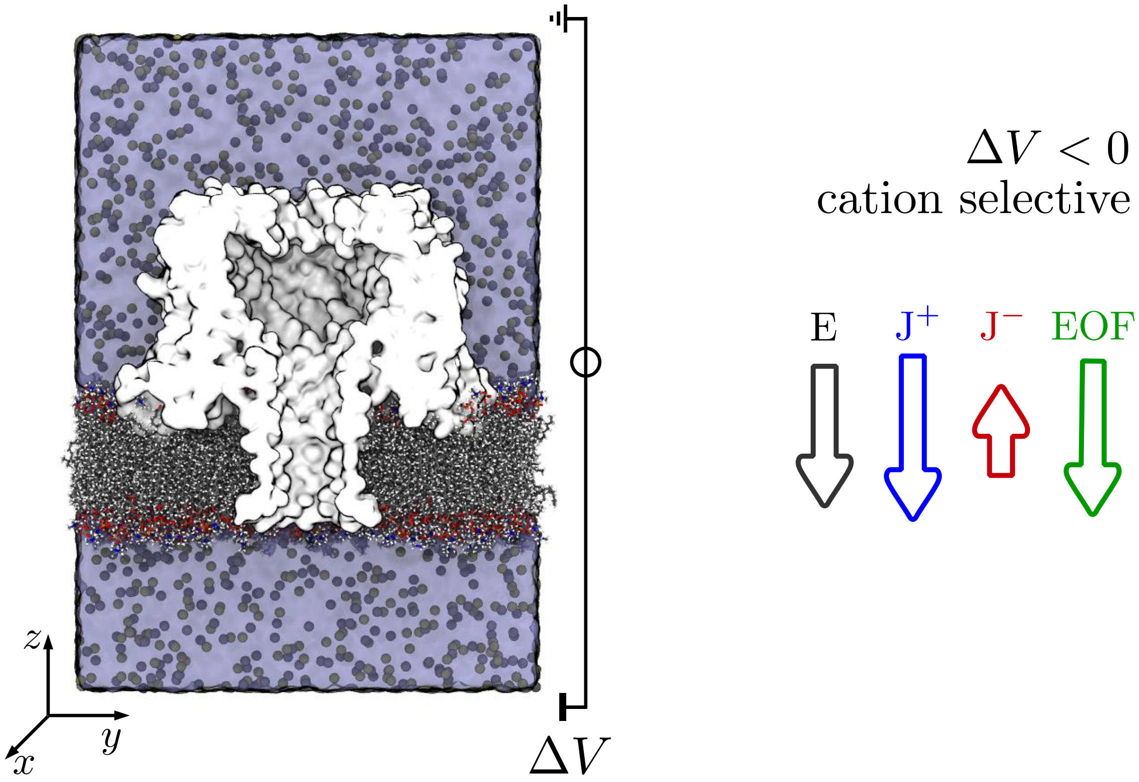

**Figure S2. MD System setup.** A triperiodic simulation box comprising a CytK pore embedded in a POPC membrane solvated with water and a 1M solution of KCl is used to study the system under different applied voltages. The system is cut along a plane parallel to the pore axis to show the pore's internal shape. Water and ions inside the pore are not represented. On the right, a sketch of the direction of the electric field, and ions and electroosmotic flow is reported for a cation selective mutant under negative voltages.

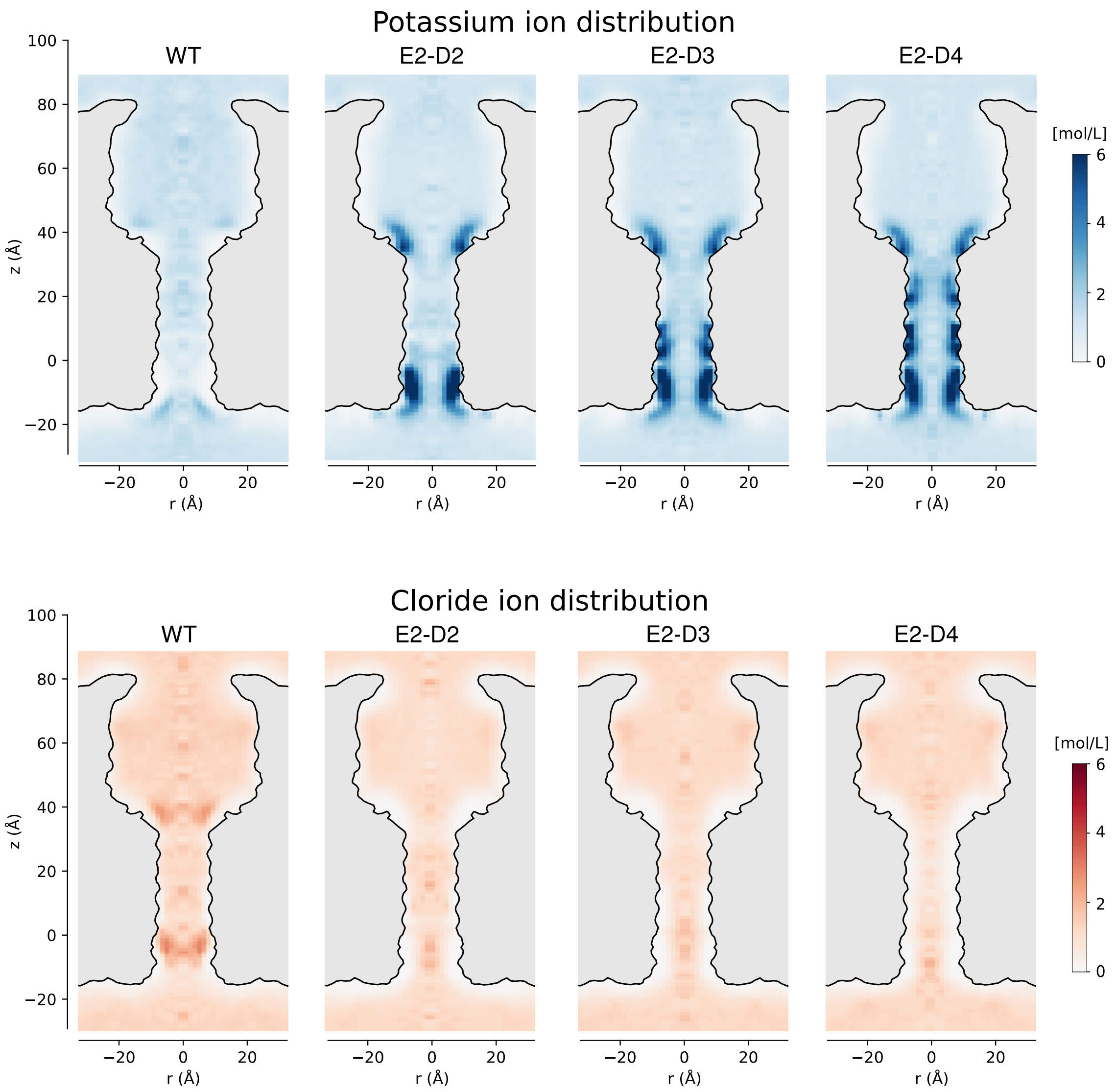

**Figure** S3. **Equilibrium maps of ionic concentration estimated by MD**. WT, K128D-K155D (2E-2D-CytK), K128D-K155D-Q145D (2E-3D-CytK) and K128D-K155D-S151D (2E-4D-CytK) pores have been simulated without applied voltage for 40 ns. Top row corresponds to $K^+$ while bottom row to $Cl^-$. Charge density is color coded with the scales on the right, expressed in molar units. The cutout of the pore is drawn to lead the eye and it roughly corresponds to the region inaccessible to the solvent. Corresponding ionic charge maps are reported in Fig 1G.

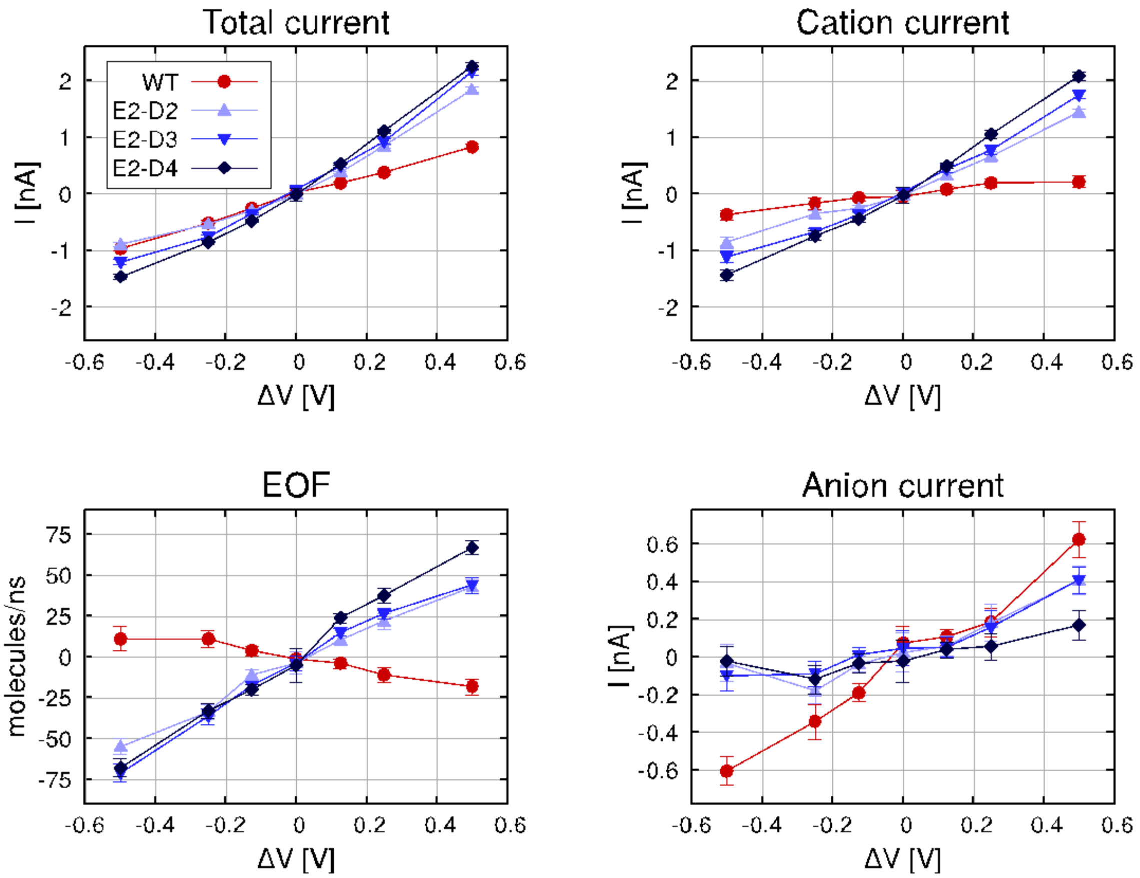

**Figure** S4. **Currents and EOF at different applied voltages estimated by MD**. The four mutants are color coded in the legend. WT is weakly anion selective, thus exhibiting a weak EOF. The mutant pores become cation selective, with selectivity increasing with every added negatively charged residue. The associated EOF is reversed with respect to WT. In all cases EOF is roughly linear in voltage, confirming the role of fixed charges in electroosmosis, and ruling out possible induced charge effects [10.1021/acsnano.1c03017].

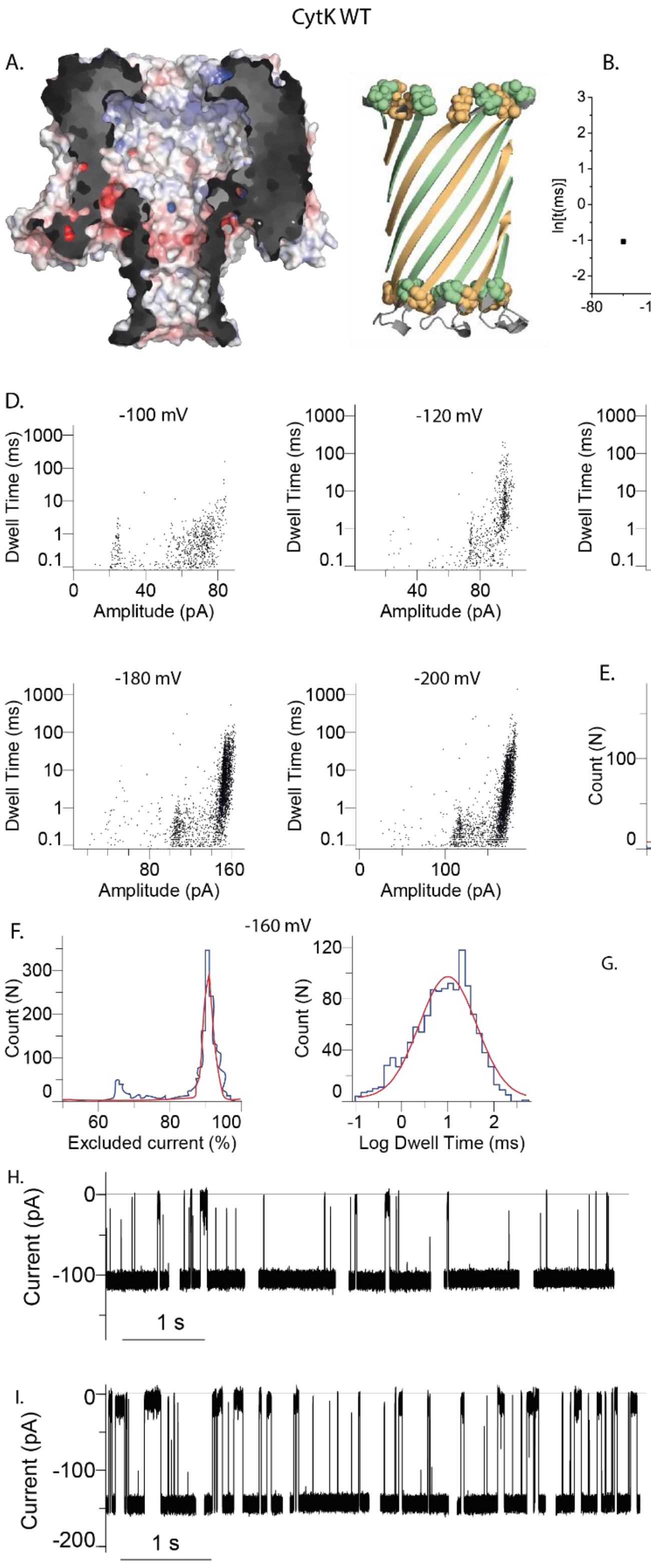

**Figure S5. S1 translocation through the CytK WT nanopore. A)** WT CytK nanopore as electrostatic surface (1 M KCl at pH 7.5) (on the left side) which shows a fairly neutral barrel and cartoon representation of the barrel (on the right), where the two pairs of charged amino acids are highlighted: E112 (green) – K155 (orange) and E139 (orange) – K128 (green). **B**) Voltage dependency of the natural logarithm of the dwell time. Two types of events were identified, a relatively long event, designated as E1, and a much shorter event cluster, E2. While E2 duration is hovering around a lnt of -1.5 to -2, the duration of E1 first increases between -100 and -140 mV, then it starts decreasing between -160 and -200 mV. **C)** $I_{ex\%}$ dependency of S1 blockades on the applied bias. **D)** Scatter plots (dwell time vs amplitude) associated with S1 capture and translocation. **E-F)** Examples of histograms obtained for the dwell time and $I_{ex}$ at -120 mV (E) and -160 mV (F). **G)** The protein sequence of S1, with positively charged residues highlighted in red. **H-I)** Typical examples of S1 capture events from at -120 mV (H) and at -160 mV (I). Recordings were carried out in 1 M KCl, 15 mM HEPES, pH 7.5.

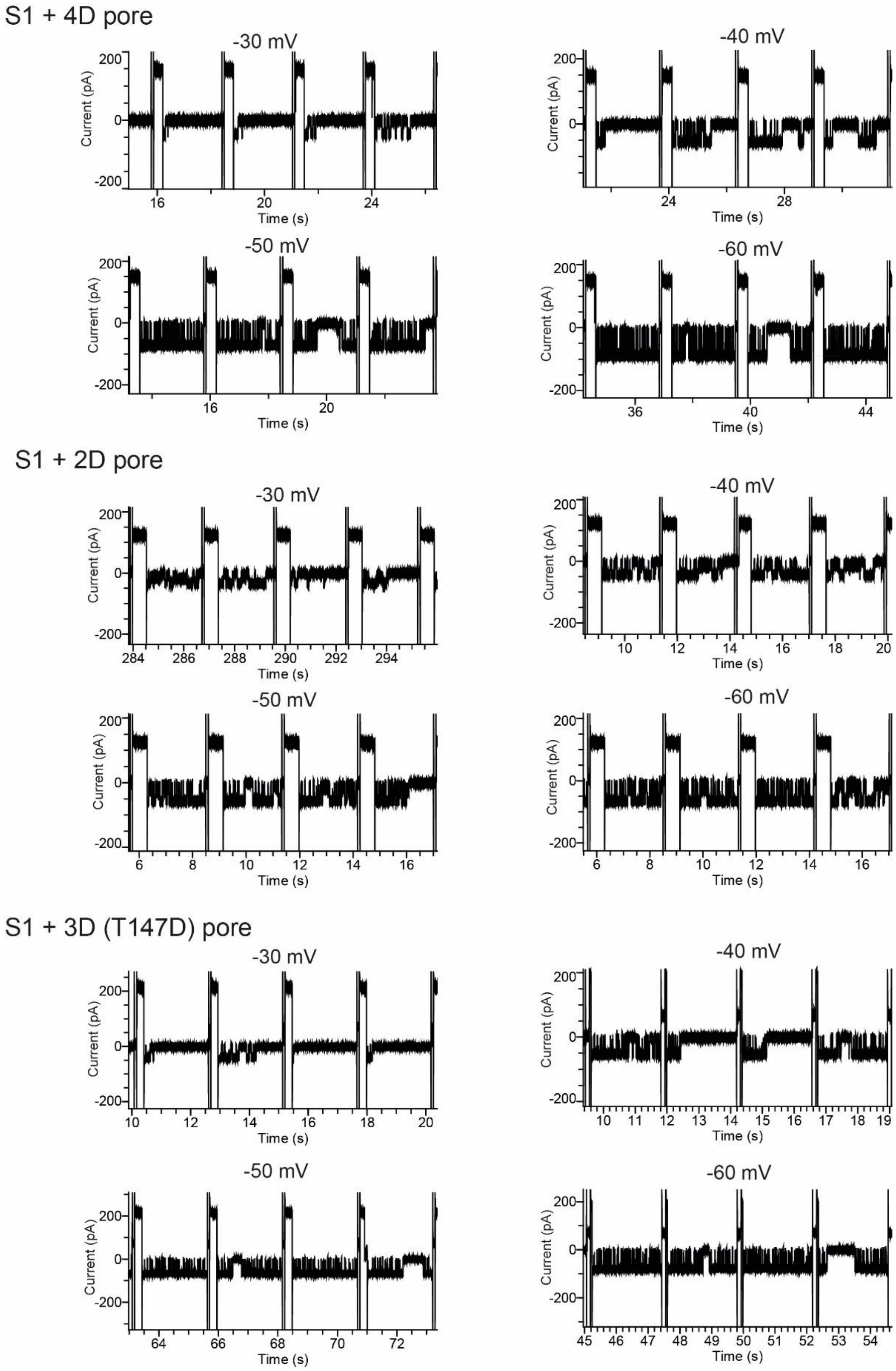

**Figure S6. S1-nanopore sticking at low potentials.** In the pores with a highly negatively charged lumen (2E-4D-CytK top, 2E-2D-CytK middle, 2E-2D-T147D-CytK bottom), the positively charged S1 may stick to the lumen at low potential, resulting in the pore being closed most of the time, while at higher potential this interaction is easier to break, promoting substrate release and thus, an open pore.

A.

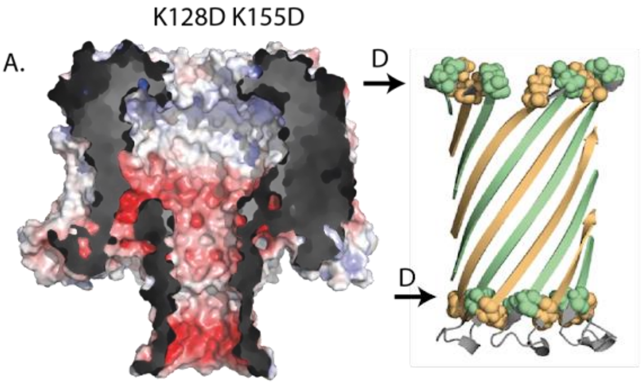

B.

S1
MGHHHHHHSSRRRRRRRRRRSSSSSSSSSSSSSSSFGYGW
SSSSSSSSSSSSSSSRRRRRRRRRRSSSSSSSSSSSSSSSFGYGW
SSSSSSSSSSSSSSSRRRRRRRRRRSSAANDENYALAA

C.

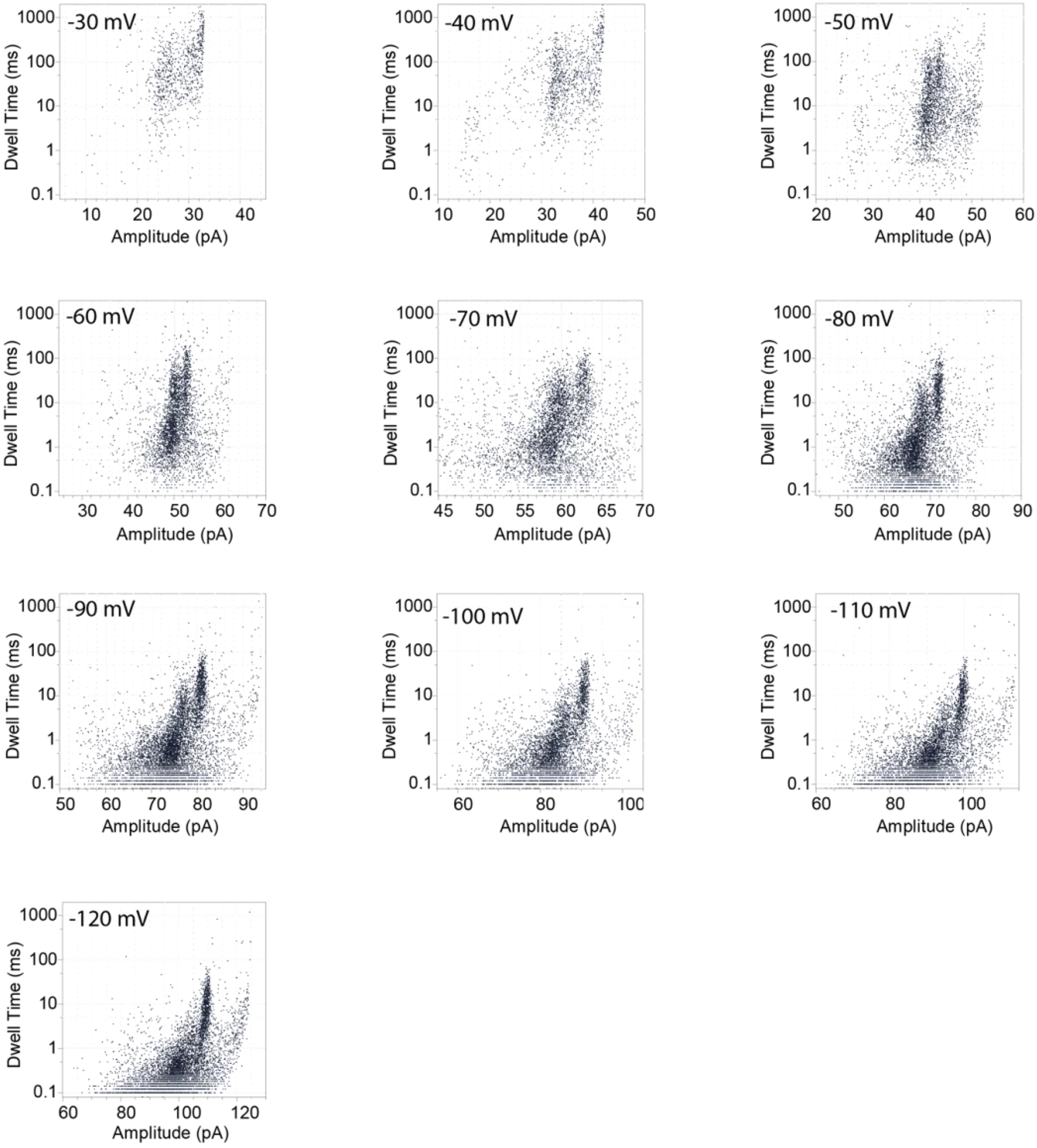

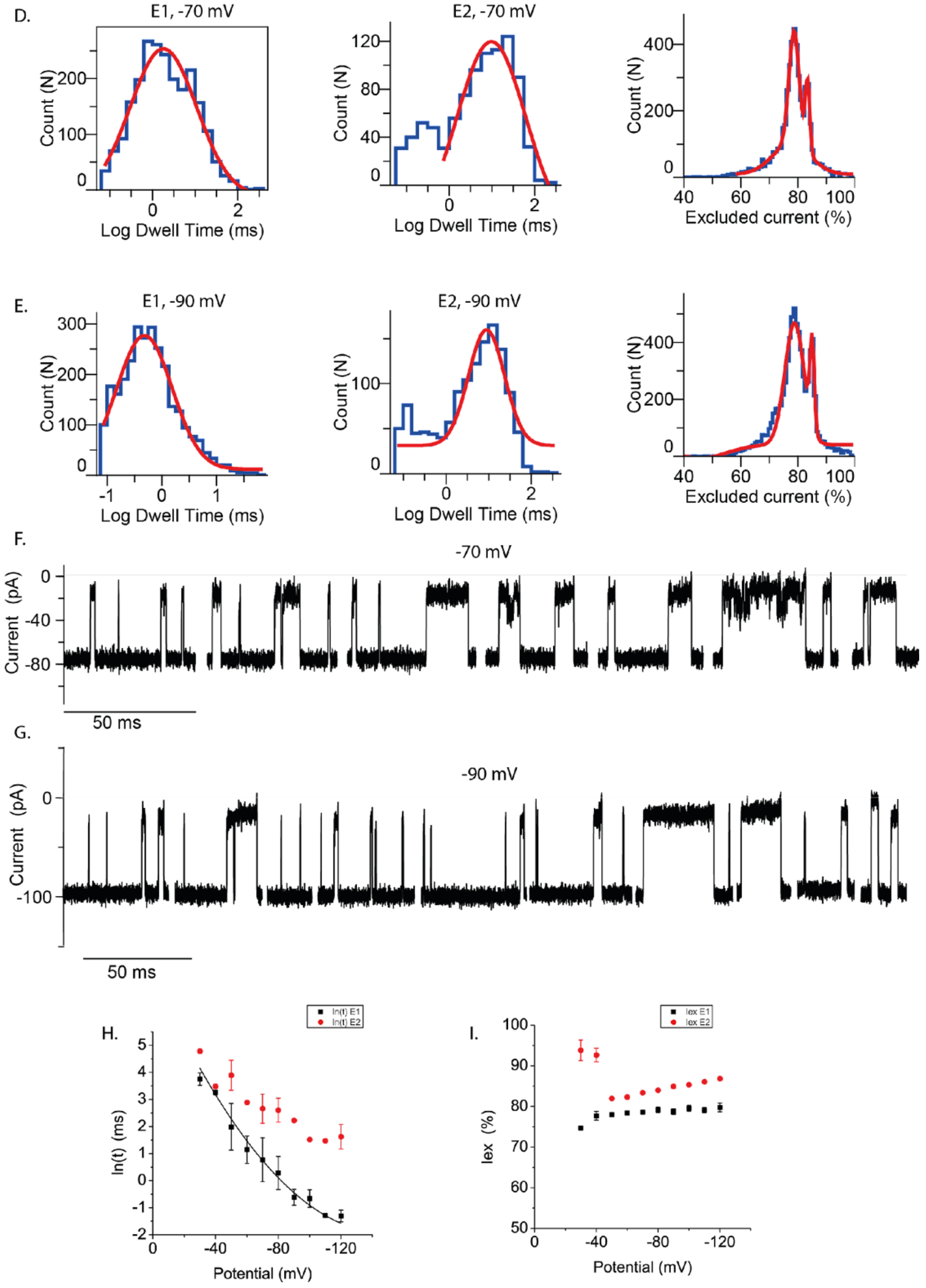

**Figure S7. S1 translocation through the K128D K155D (2E-2D) CytK mutant nanopore. A)** Electrostatic surface representation (1 M KCl at pH 7.5) of K128D K155DCytK (2E-2D-CytK),left; and cartoon representation of the transmembrane barrel, right. The replacement of the positive charges with negative charges renders the extremities of the barrel relatively negatively charged at physiological pH. In the cartoon representation only the negatively charged residues are shown as spheres: E112 (green), E139 (orange) and K128D (green). **B)** The amino acid sequence of the S1 substrate, with the positively charged residues shown in red and the acid ones in blue. **C)** A set of scatter plots from the investigated voltages. **D-E)** Examples of histograms obtained for the dwell time and $I_{ex}$ at -70 mV (D) and -90 mV (E). **F-G)** Typical examples of S1 capture events froM at -70 mV (F) and at -90 mV (G). Recordings were carried out in 1 M KCl, 15 mM HEPES, pH 7.5. **H)** Voltage dependency of the natural logarithm of the dwell time. Two types of events were identified, E1 and E2. **I)** $I_{ex\%}$ dependency of the E1 and E2 S1 blockades on the applied bias.

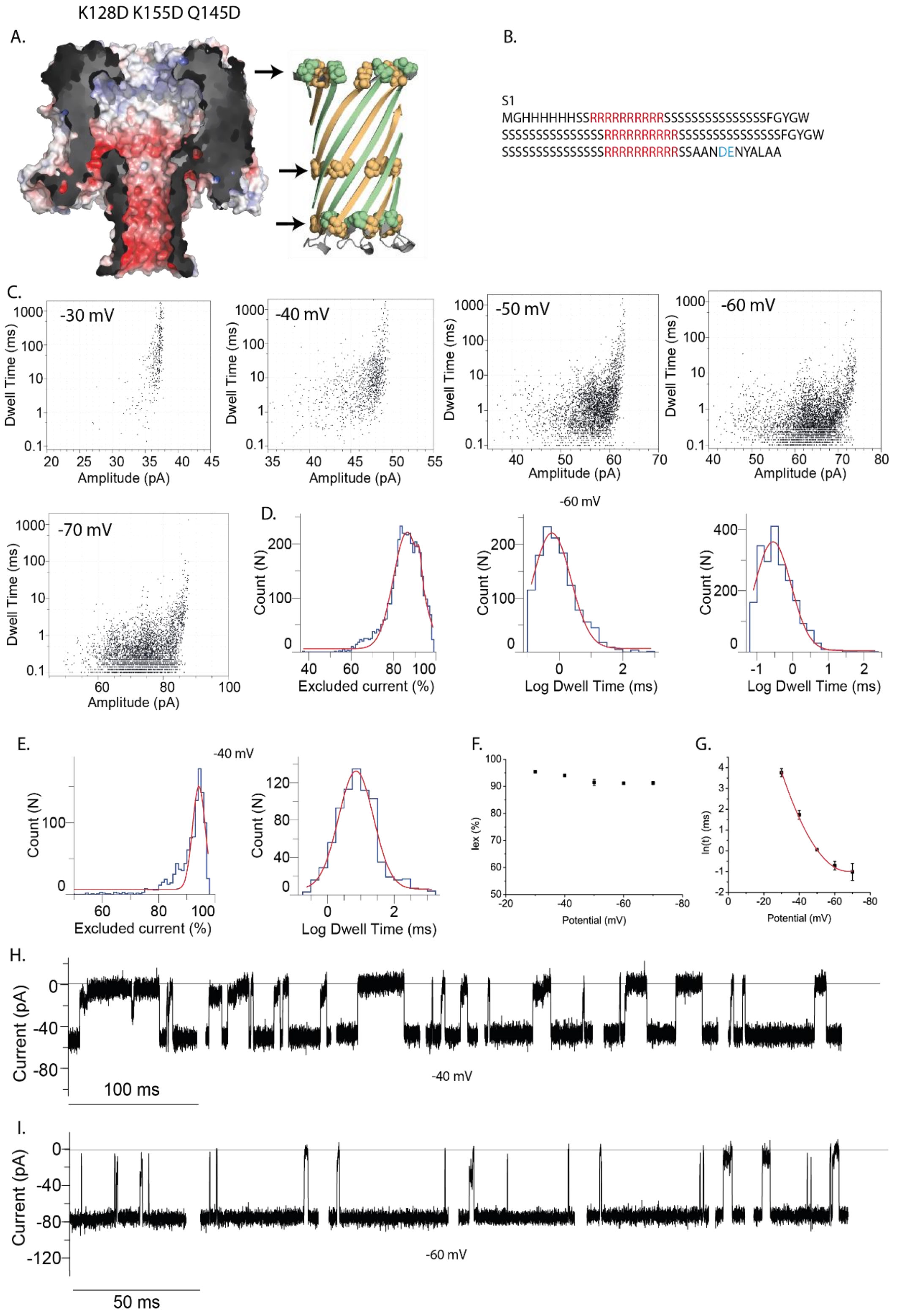

**Figure S8. S1 translocation through the K128D K155D Q145D (2E-2D-Q145D) CytK mutant nanopore**. **A)** Depiction of the 2E-2D-Q145D-CytK nanopore as electrostatic surface (1 M KCl at pH 7.5), left and cartoon representation (mutated residues shown as spheres), right. **B)** S1 protein sequence with positively charged residues highlighted in red. **C)** Scatter plots (dwell time vs amplitude) associated with S1 capture and translocation. **D-E)** Examples of histograms obtained for the dwell time and $I_{ex}$ at -60 mV (D) and -40 mV (E). **F)** Voltage dependency of the natural logarithm of the dwell time. **G)** Voltage dependency of the natural logarithm of the dwell time. **H-I)** Typical examples of S1 capture events froM at -40 mV (H) and at -60 mV (I). Recordings were carried out in 1 M KCl, 15 mM HEPES, pH 7.5.

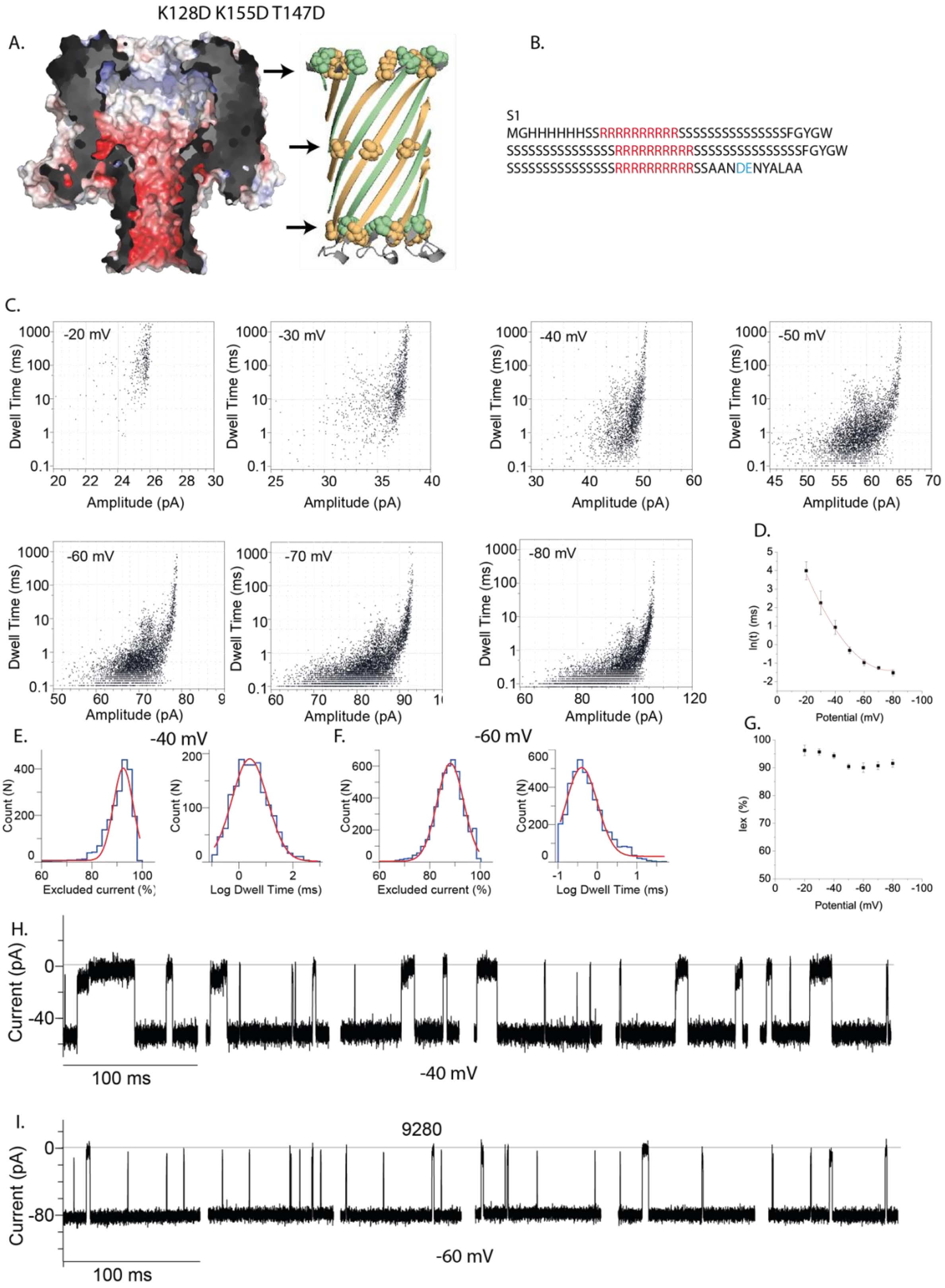

**Figure S9. S1 translocation through the K128D K155D T147D (2E-2D-T147D) CytK mutant nanopore. A)** Illustration of the CytK K128D K155D T147D (2E-2D-T147D-CytK) nanopore as electrostatic surface (1 M KCl at pH 7.5), left; and cartoon representation of the barrel, right, where the substitutions are highlighted. **B**) Sequence of S1. **C)** Scatter plots (dwell time vs amplitude) associated with S1 capture and translocation events. **D)** Voltage dependency of the natural logarithm of the dwell time. E1 and E2 events become indistinguishable. The dwell time is calculated from all the events. **E-F)** Examples of histograms obtained for the dwell time and $I_{ex}$ at -40 mV (E) and -60 mV (F). **G)** $I_{ex\%}$ dependency of S1 blockades on the applied bias. **H-I)** Typical examples of S1 capture events from at -40 mV (H) and at -60 mV (I). Recordings were carried out in 1 M KCl, 15 mM HEPES, pH 7.5.

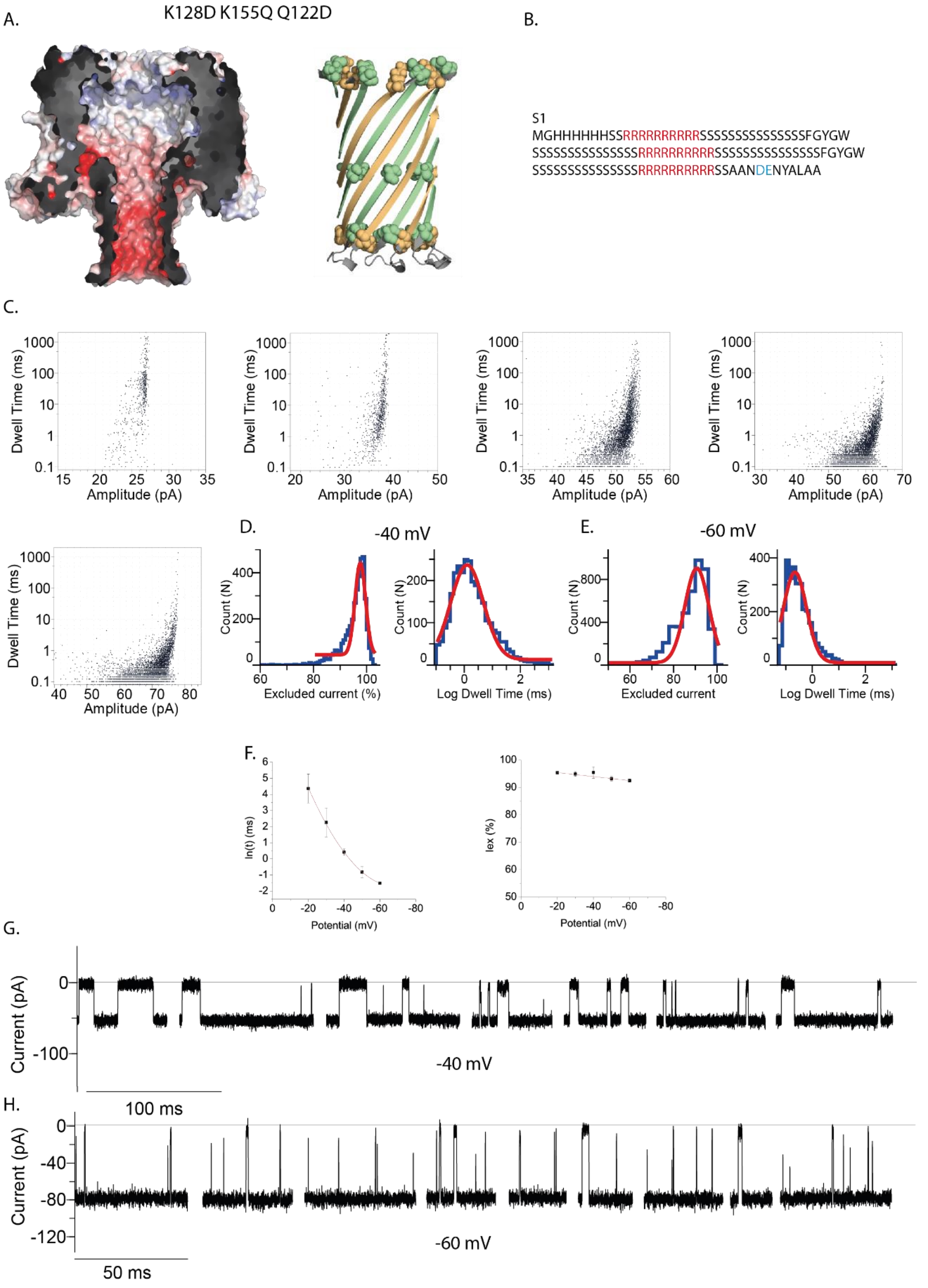

**Figure S10. Translocation of S1 through the 2E-1D-1Q-Q122D-CytK nanopore**. **A)** Illustration of the CytK-K128D-K155Q-Q122D (2E-1D-1Q-Q122D-CytK) nanopore as electrostatic surface (1 M KCl at pH 7.5), left; and cartoon representation of the barrel, right, where the substitutions are highlighted. **B**) S1 protein sequence. **C)** Scatter plots (dwell time vs amplitude) associated with S1 capture and translocation. **D-E)** Examples of histograms obtained for the dwell time and $I_{ex}$ at -40 mV (D) and -60 mV (E). **F)** Voltage dependency of the natural logarithm of the dwell time, left; and $I_{ex\%}$ dependency of S1 blockades on the applied bias, right. **G-H)** Typical examples of S1 capture events from at -40 mV (G) and at -60 mV (H). Recordings were carried out in 1 M KCl, 15 mM HEPES, pH 7.5.

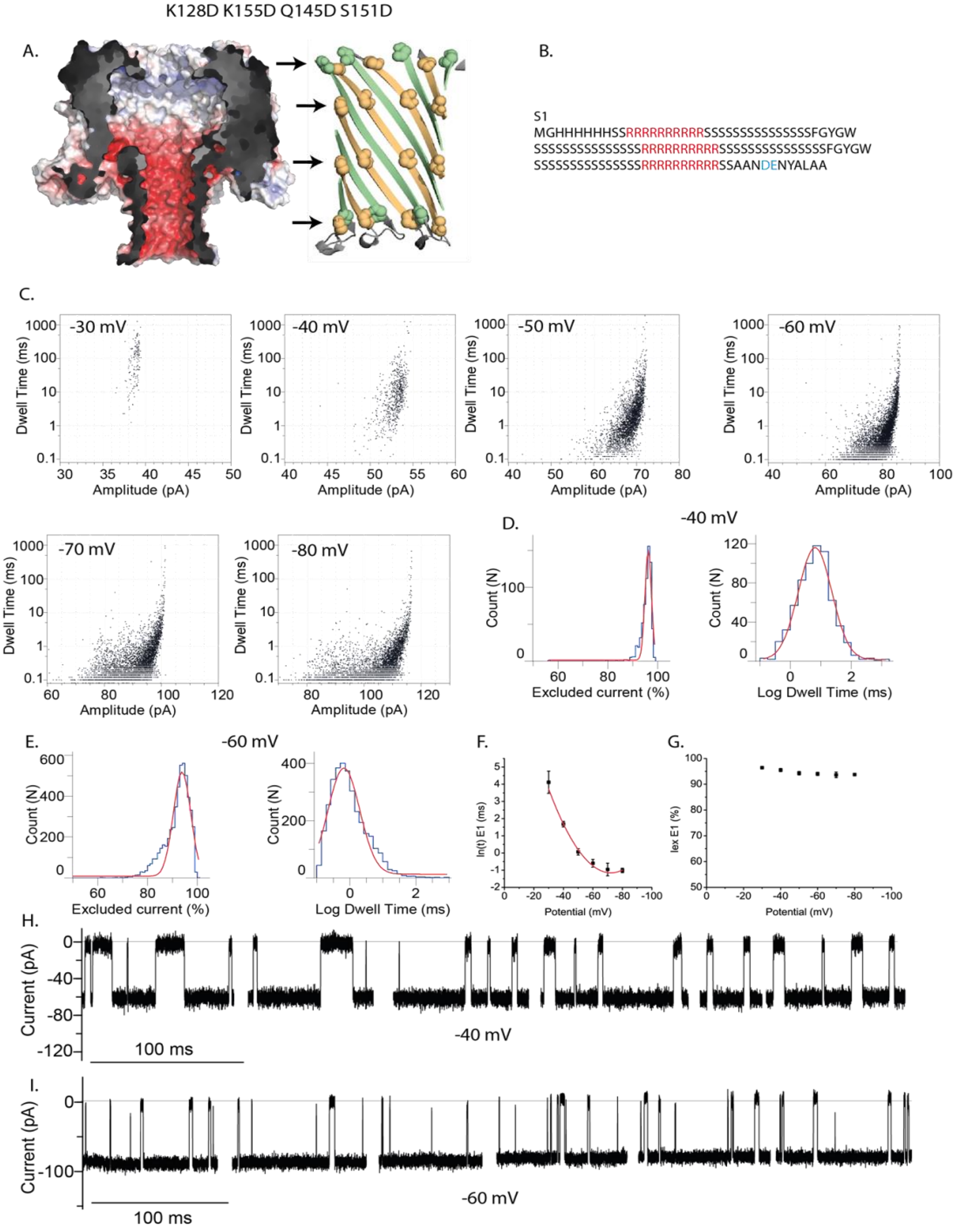

**Figure S11. Translocation of S1 through CytK-2E-4D nanopores**. **A)** Electrostatic surface illustration of the K128D-K155D-Q145D-S151D-CytK (2E-4D-CytK) nanopore in 1 M KCl at pH 7.5, left; and cartoon representation of the barrel where the substitutions are highlighted, right. **B**) The protein sequence of S1, with positively charged residues highlighted in red. **C)** Scatter plots (dwell time vs amplitude) associated with S1 capture and translocation. **D-E)** Examples of histograms obtained for the dwell time and $I_{ex}$ at -40 mV (D) and -60 mV (E). **F)** Voltage dependency of the natural logarithm of the dwell time. **G)** $I_{ex\%}$ dependency of S1 blockades on the applied bias. **H-I)** Typical examples of S1 capture events from at -40 mV (H) and at -60 mV (I). Recordings were carried out in 1 M KCl, 15 mM HEPES, pH 7.5.

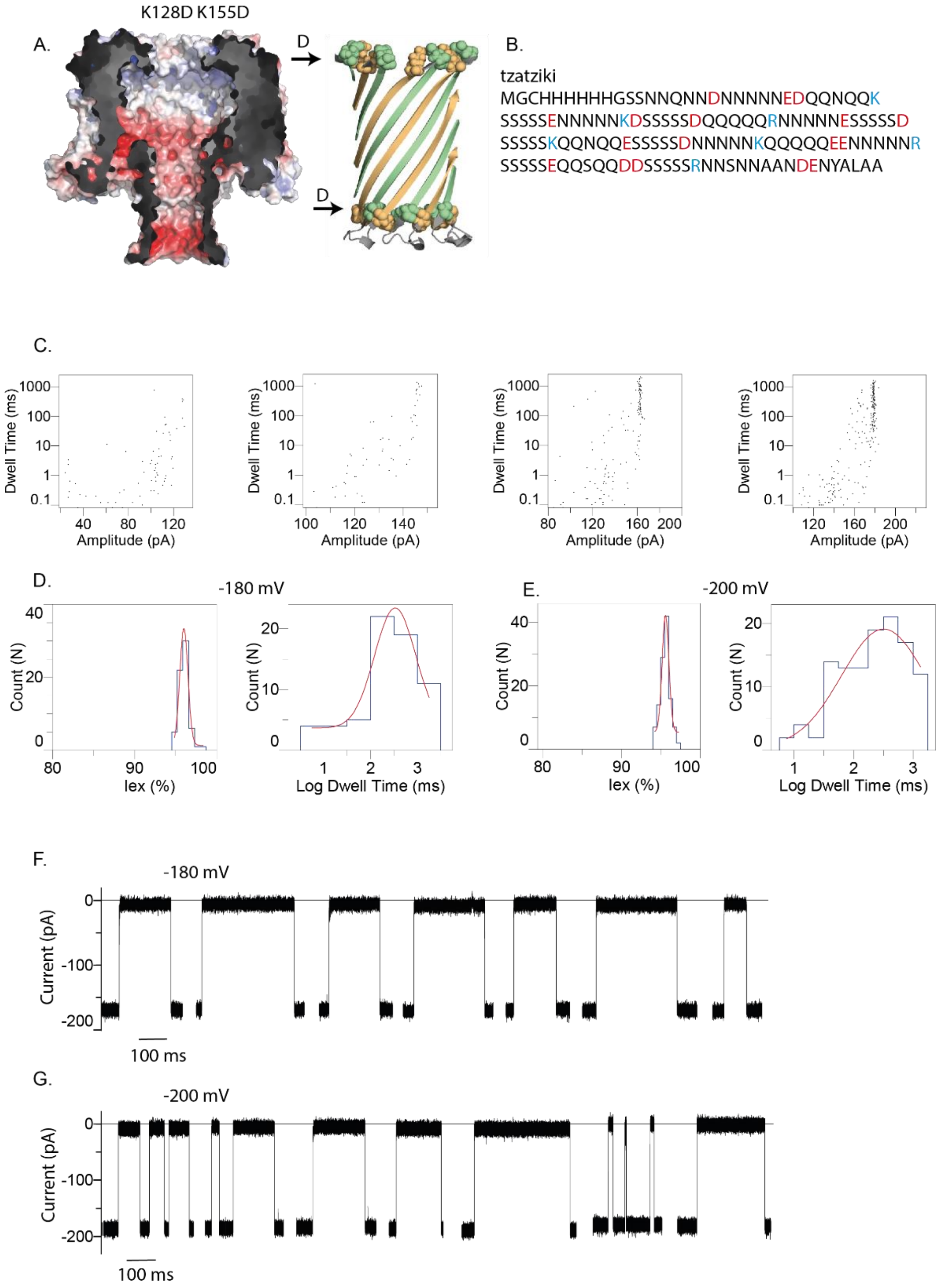

**Figure S12. Tzatziki and the CytK 2E-2D nanopore**. **A)** Electrostatic surface illustration of K128D-K155D-CytK (2E-2D-CytK) nanopore in 1 M KCl at pH 7.5, left; and cartoon representation of the barrel where the substitutions are highlighted, right. **B**) The protein sequence of tzatziki, with positively charged residues highlighted in red and negatively charged residues in blue. **C)** Scatter plots (dwell time vs amplitude) associated with tzatziki capture at -140, -160, -180 and -200 mV. The first two potentials are associated with sporadic events, which do not seem to form a cluster. **D-E)** Examples of histograms obtained for the dwell time and $I_{ex}$ at -180 mV (D) and -200 mV (E). **F-G)** Typical examples of tzatziki capture events from at -180 mV (H) and at -200 mV (I). Recordings were carried out in 1 M KCl, 15 mM HEPES, pH 7.5.

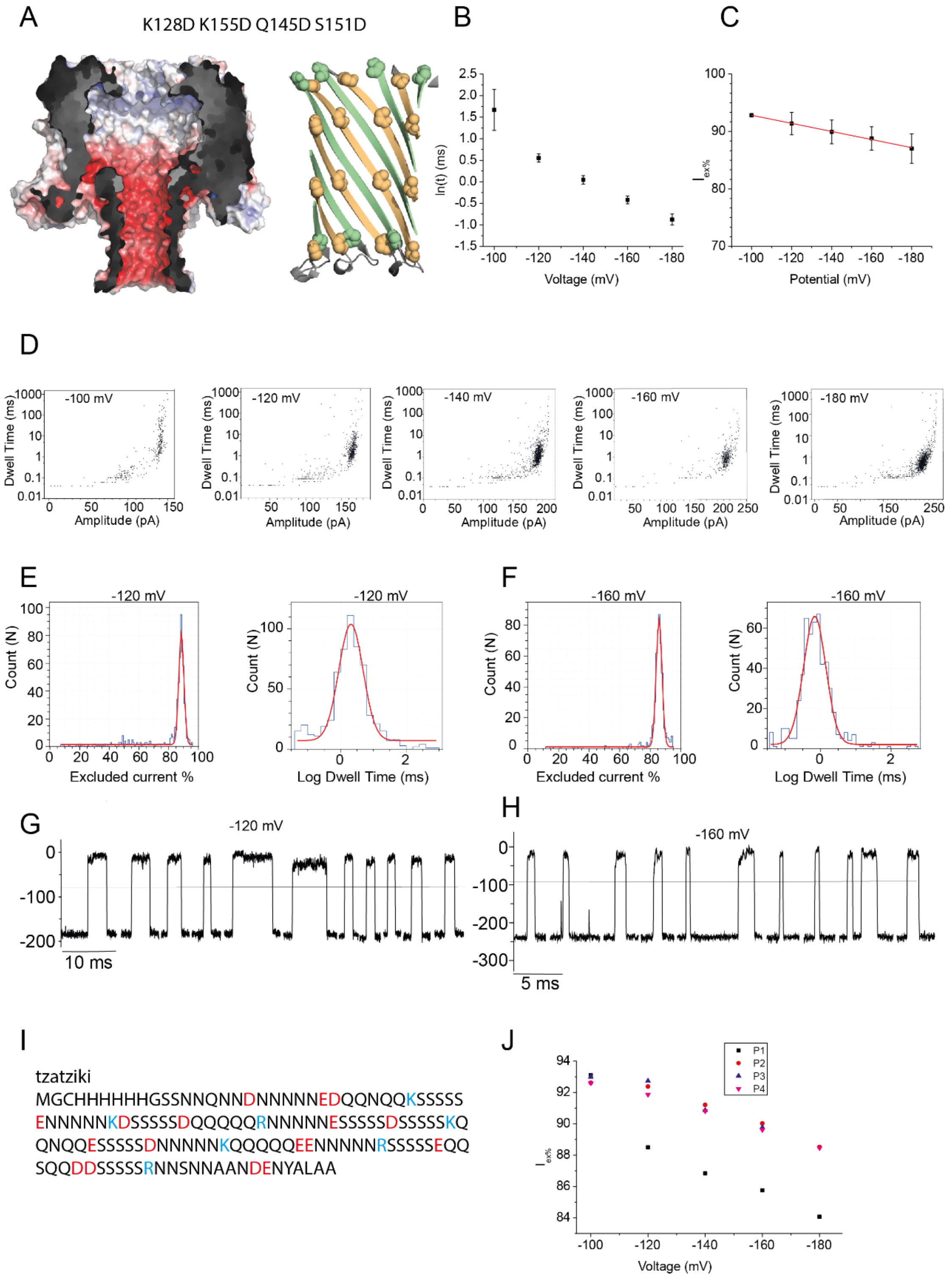

**Figure S13. Tzatziki translocation through the K128D Q145D S151D K155D (2E-4D) CytK nanopore. A)** Electrostatic surface illustration of the K128D-K155D-Q145D-S151D-CytK (2E-4D-CytK) nanopore in 1 M KCl at pH 7.5, left; and cartoon representation of the barrel where the substitutions are highlighted, right. **B**) Voltage dependency of the natural logarithm of the dwell time. **C)** $I_{ex\%}$ dependency of tzatziki blockades on the applied bias. **D)** Scatter plots (dwell time vs amplitude) associated with tzatziki capture and translocation. **E-F)** Examples of histograms obtained for the dwell time and $I_{ex}$ at -120 mV (E) and -160 mV (F). **G-H)** Typical examples of tzatziki capture events from at -120 mV (G) and at -160 mV (H). **I)** Protein sequence of tzatziki, with positively charged residues highlighted in red and negatively charged residues in blue. Recordings were carried out in 1 M KCl, 15 mM HEPES, pH 7.5.

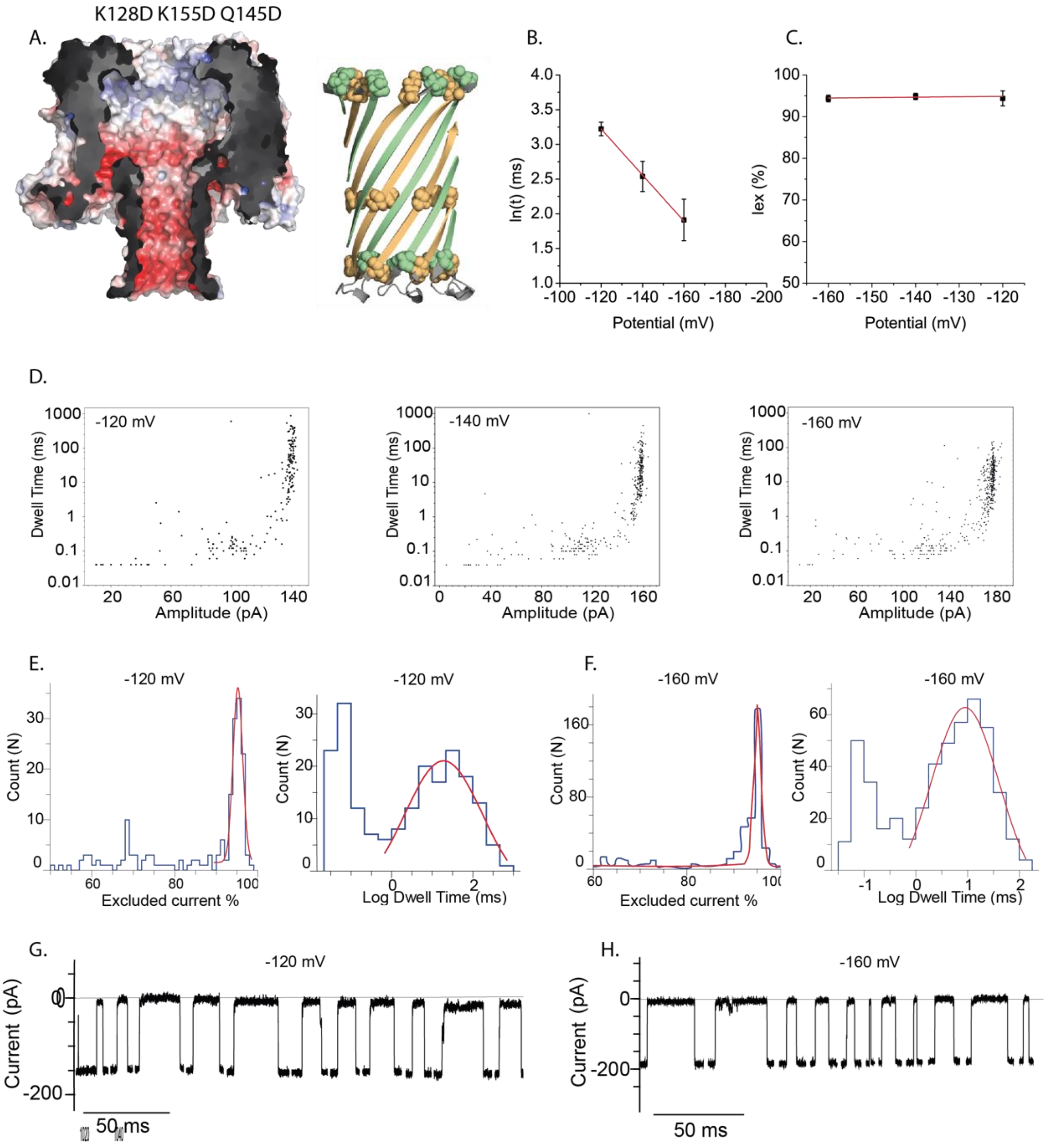

I.

tzatziki
MGCHHHHHHGSSNNQNNDNNNNNEDQQNQQKSSSSSENNNNNKDSSSSSDQQQQQRNNNNNESSSSSD
SSSSSKQQNQQESSSSSDNNNNNKQQQQQEENNNNNRSSSSSEQQSQQDDSSSSSRNNSNNAANDENYALAA

**Figure S14. Tzatziki translocation through the K128D K155D Q145D (2E-2D-Q145D) CytK nanopore. . A)** Electrostatic surface illustration of the K128D-K155D-Q145D-CytK (2E-2D-Q145D-CytK) nanopore in 1 M KCl at pH 7.5, left; and cartoon representation of the barrel where the substitutions are highlighted, right. **B**) Voltage dependency of the natural logarithm of the dwell time. **C)** $I_{ex\%}$ dependency of tzatziki blockades on the applied bias. **D)** Scatter plots (dwell time vs amplitude) associated with tzatziki capture and translocation. **E-F)** Examples of histograms obtained for the dwell time and $I_{ex}$ at -120 mV (E) and -160 mV (F). **G-H)** Typical examples of tzatziki capture events from at -120 mV (G) and at -160 mV (H). Recordings were carried out in 1 M KCl, 15 mM HEPES, pH 7.5. **I)** The protein sequence of tzatziki, with positively charged residues highlighted in red and negatively charged residues in blue.

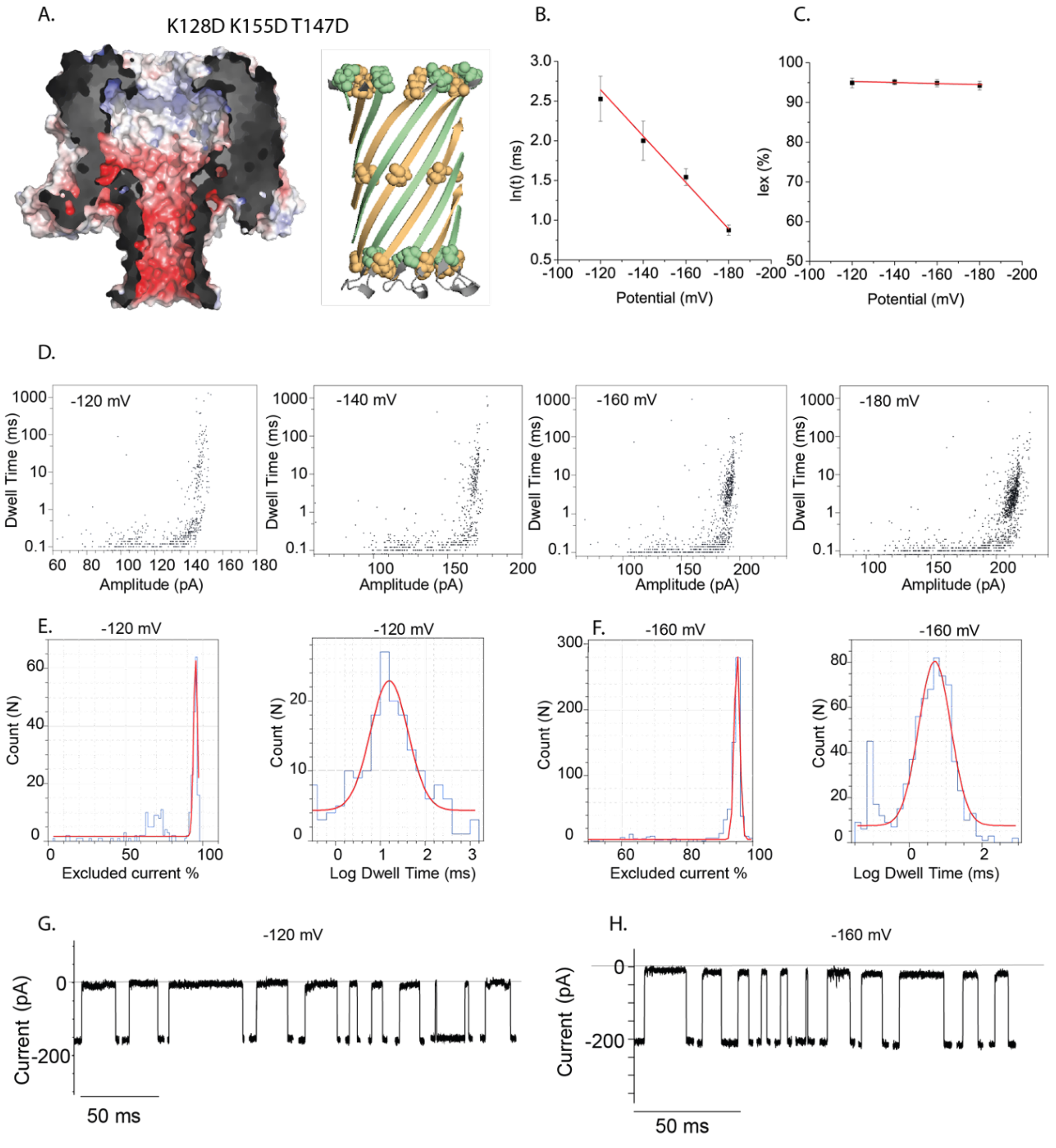

I.

tzatziki
MGCHHHHHHGSSNNQNNDNNNNNEDQQNQQKSSSSSENNNNNKDSSSSSDQQQQQRNNNNNESSSSSD
SSSSSKQQNQQESSSSSDNNNNNKQQQQQEENNNNNRSSSSSEQQSQQDDSSSSSRNNSNNAANDENYALAA

**Figure S15. Tzatziki translocation through the K128D K155D T147D (2E-2D-T147D) CytK mutant nanopore. A)** Electrostatic surface illustration of the K128D-K155D-T147D-CytK (2E-2D-T147D-CytK) nanopore in 1 M KCl at pH 7.5, left; and cartoon representation of the barrel where the substitutions are highlighted, right. **B**) Voltage dependency of the natural logarithm of the dwell time. **C)** $I_{ex\%}$ dependency of tzatziki blockades on the applied bias. **D)** Scatter plots (dwell time vs amplitude) associated with tzatziki capture and translocation. **E-F)** Examples of histograms obtained for the dwell time and $I_{ex}$ at -120 mV (E) and -160 mV (F). **G-H)** Typical examples of tzatziki capture events from at -120 mV (G) and at -160 mV (H). Recordings were carried out in 1 M KCl, 15 mM HEPES, pH 7.5. **I)** The protein sequence of tzatziki, with positively charged residues highlighted in red and negatively charged residues in blue.

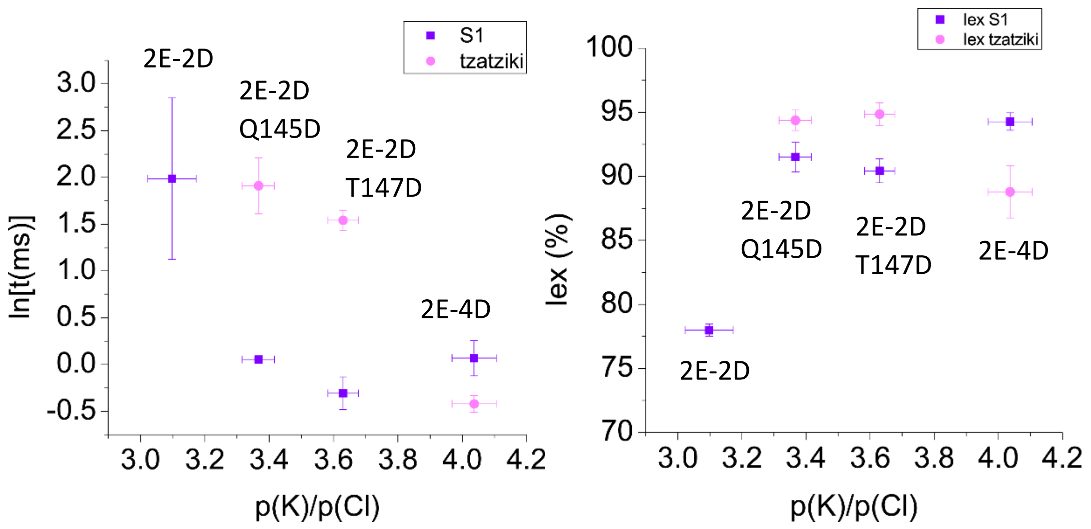

**Figure S16. Cation selectivity vs ln(dwell time), left; and $I_{ex\%}$, right.** The dwell time decreased with the strength of the EOF (related to the cation selectivity), indicating the EOF drives translocation. An exception to the trend was the transport of S1 through 2E-4D, which may be explained by the sticking of the positive charges of S1 to the negative charges of the nanopore. Using S1, it was observed an increased $I_{ex}$ with the number of introduced aspartate residues introduced. Most likely, as the pore becomes more cation selective, a larger proportion of cation is blocked by the positively charged substrate. This effect was not observed with tzatziki. A reduction of current with the 2E-4D-CytK respect to the other nanopore most likely reflects the increased stretching of the polypeptide. The voltage was set to -50 mV when testing S1 and at -160 mV for tzatziki.

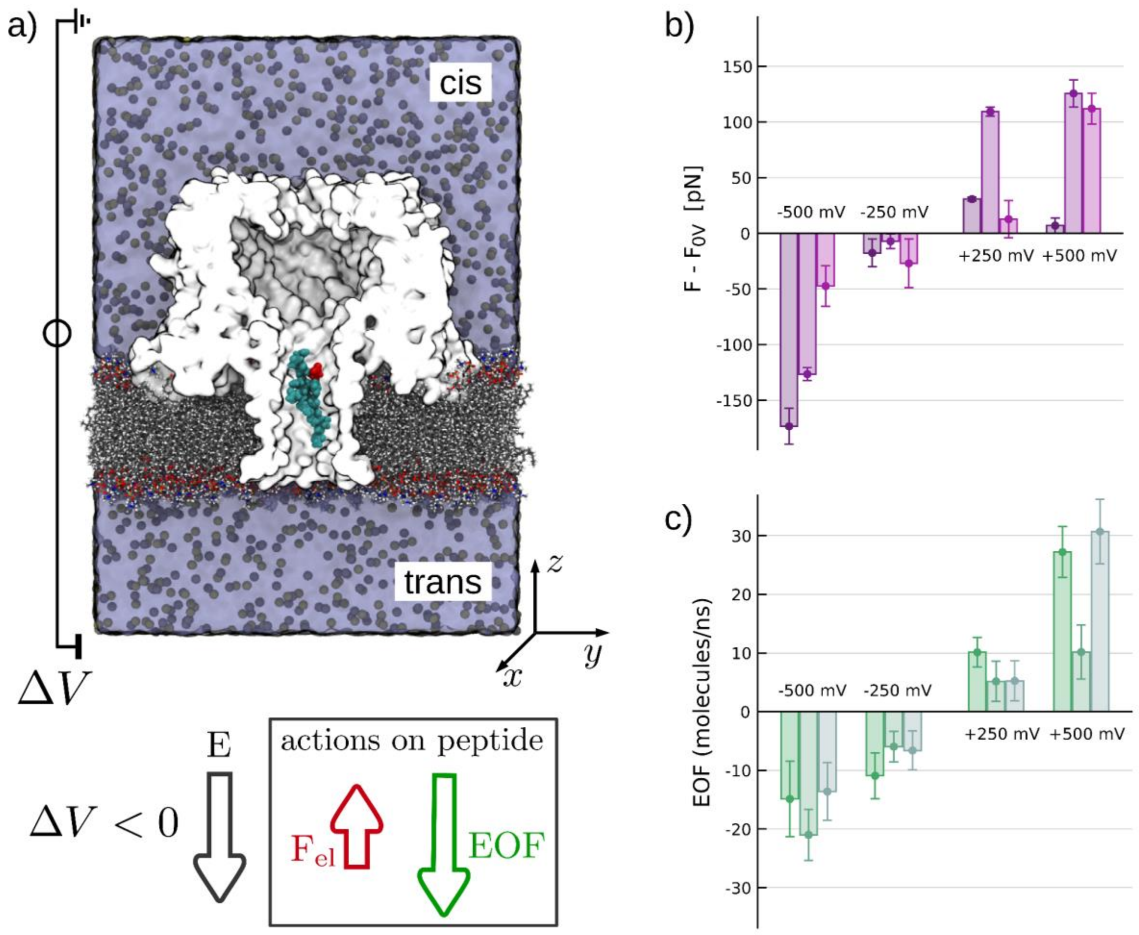

**Figure S17. Forces on a short Tzatziki fragment in the 2E-4D-Cytk barrel. a)** Sketch of the simulation set-up. The 11-mer peptide of sequence GSSNNQNNDNN, corresponding to the Tzatziki residues 10-20, is preliminary pulled in the nanopore barrel. The N-term points toward the trans side. Both terminals are neutralized, in particular, an acetyl group is used to terminate the N-term while an amine group to the C-term. Consequently, the only charged residue is the ASP, red in the sketch. The system is cut along a plane parallel to the pore axis to show the pore's internal shape. Water and ions inside the pore are not represented. The z-coordinate of the peptide center of mass is constrained to its initial position with a harmonic spring and, a voltage is applied to the system. **b)** Force on the peptide. We performed three independent replicas, each starting before the importing of the peptide. For each replica, we used the final conformation obtained after importing and equilibration as the initial state for all the production runs. Specifically, we run five production simulations at voltages -500 mV, -250 mV, 0 mV, +250 mV, +500 mV. The effect of the combined action of the electrophoretic force $F_{el}$ acting on the ASP (negative) residue and the electroosmotic flow (EOF) is quantified by the difference in the average force along the axis between the biased simulation and the 0V-case. The duration of production runs is 60 ns at +/- 250 mV and 30 ns at +/- 500 mV and 0 mV. For each voltage, the three bars of different colors correspond to the independent replicas. The corresponding EOF is reported in panel **c)**. Taken together, the resulting force is always directed as the EOF, indicating that competition between the drag on the peptide due to EOF and the electrophoresis is dominated by the EOF. For negative ΔV, depicted in the lower left corner, this would result in a cis-to-trans movement. For comparison, we also applied the same protocol to a homopeptide of 11 Arginines. In this case, EOF and electrophoresis cooperate and, coherently, the forces are much larger (~100 pN at 250 mV). Confidence intervals for each run are estimated via block average. panel a was made using VMD.[12]

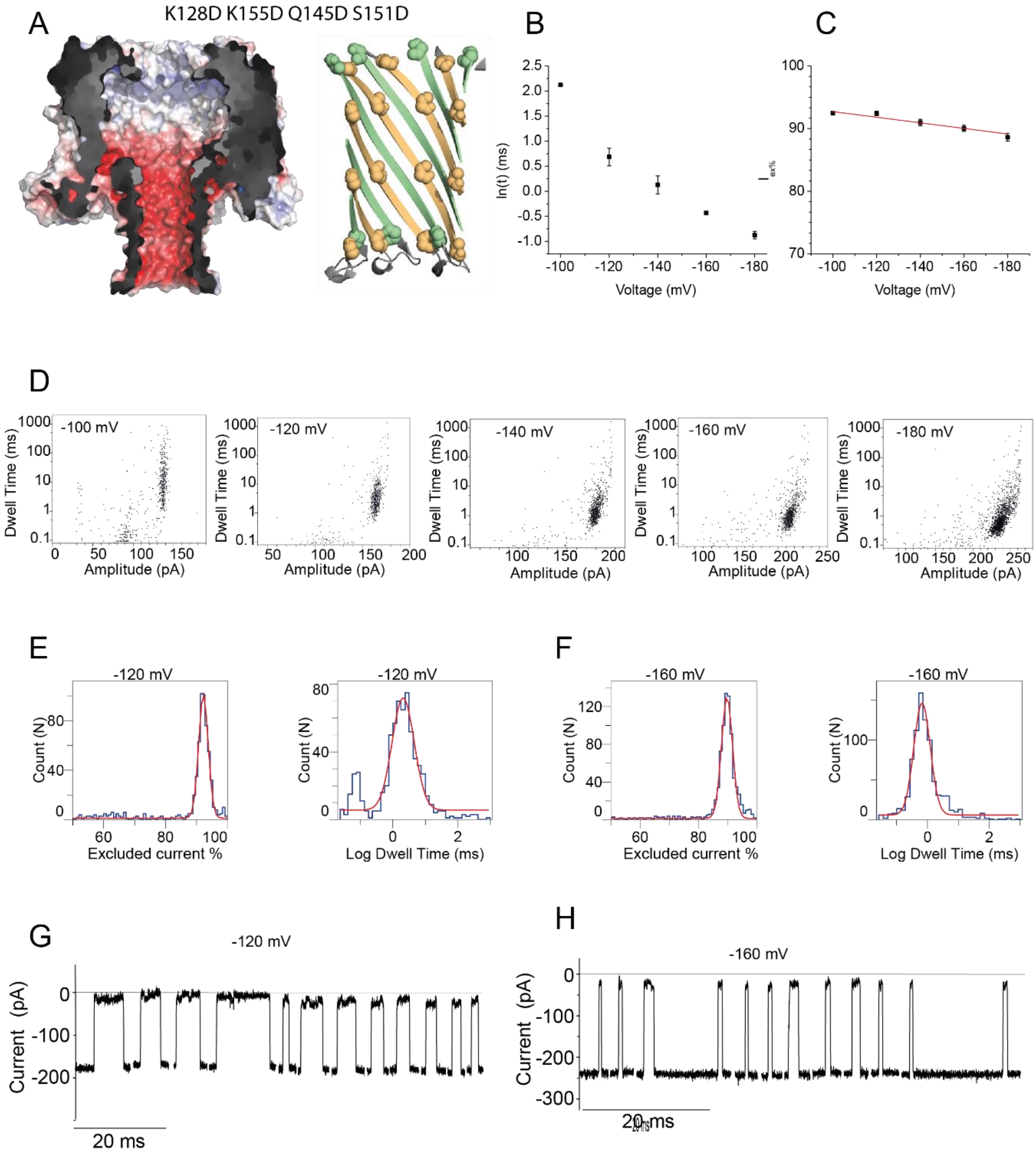

I

mujdei
MGCHHHHHHGSSNNQNNDNNNNNEDQQNQQKSSSSSENNNNNKDSSSSSDQQQQQRNNNNNESSSSSD
SSSSSKQEDEEESSSSSDNNNNNKQQQQQEENNNNNRSSSSSEQQSQQDDSSSSSRNNSNNAANDENYALAA

**Figure S18. Translocation of mujdei through CytK-2E-4D nanopores**. **A)** Electrostatic surface illustration of the K128D-K155D-Q145D-S151D-CytK (2E-4D-CytK) nanopore in 1 M KCl at pH 7.5, left; and cartoon representation of the barrel where the substitutions are highlighted, right. **B**) Voltage dependency of the natural logarithm of the dwell time. **C)** $I_{ex\%}$ dependency of mujdei blockades on the applied bias. **D)** Scatter plots (dwell time vs amplitude) associated with tzatziki capture and translocation. **E-F)** Examples of histograms obtained for the dwell time and $I_{ex}$ at -120 mV (E) and -160 mV (F). **G-H)** Typical examples of mujdei capture events from at -120 mV (G) and at -160 mV (H). Recordings were carried out in 1 M KCl, 15 mM HEPES, pH 7.5. **I)** The protein sequence of mujdei, with positively charged residues highlighted in red and negatively charged residues in blue.

*Positively charged barrel nanopore mutants*

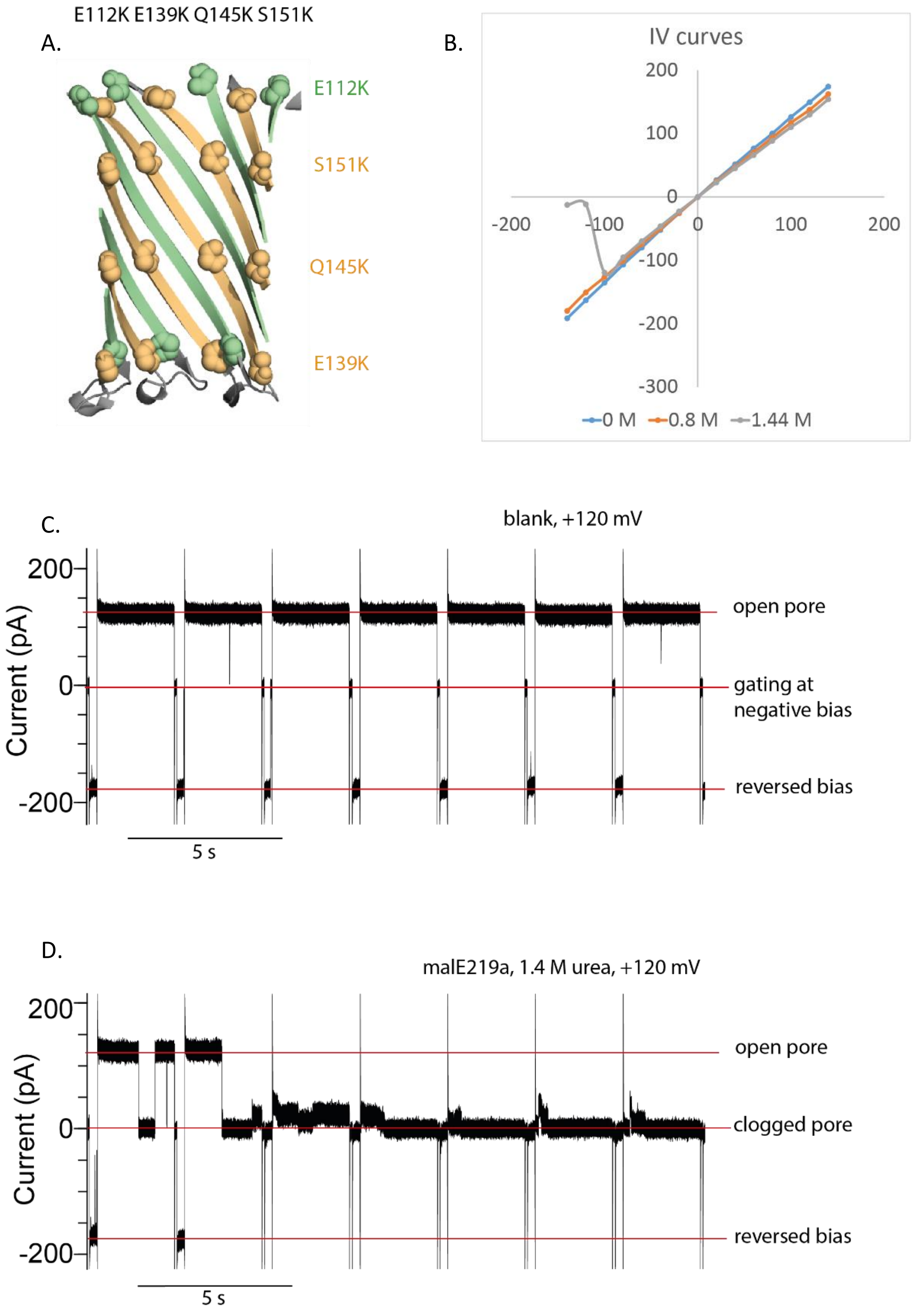

**Figure S19. The designed CytK 6K nanopore. A)** Cartoon depiction of the barrel of the CytK nanopore. Six residues were introduced were substituted to lysine, E112K, E139K, Q145K and S151K, which are illustrated as spheres. This pore is the lysine analogue of the 2E-4D pore, as the. **B)** I-V curves from the CytK 6K mutant under different concentrations of urea. Note that this mutant gated quite significantly at negative voltage, which is why the I-V curve slope is disrupted at the high negative potentials in the case of the 1.4 M urea curve. **C)** Typical current of a blank measurement in 1.4 M urea showing the pore is mostly open at positive applied bias (+120 mV). When the bias is reversed to -150 mV, the pore spontaneously closed (gating). **D)** The addition of the malE219a protein (*cis*) resulted in the pore being almost irreversibly clogged, due to the highly unlikely release of the substrate. In the unlikely situation that the pore could be opened, it would be immediately clogged by the substrate and any attempts to measure anything else than clogging were futile. Recordings were performed in 1 M KCl, 15 mM HEPES, 1.4 M urea, pH 7.5.

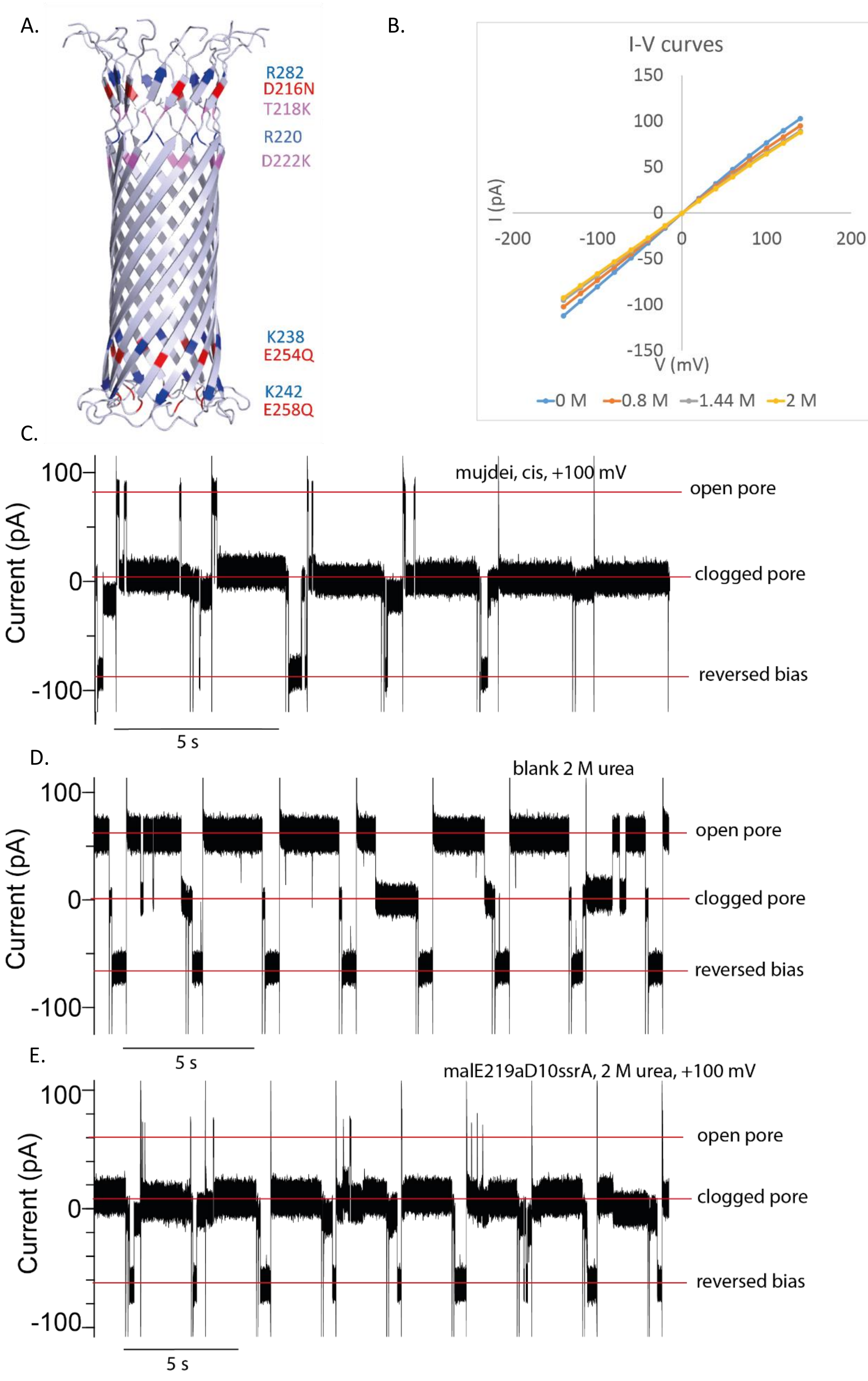

**Figure S20. Aerolysin mutant with positive charges introduced in the barrel. A)** The barrel of the aerolysin nanopore is depicted together with the modifications that resulted in the mutant designated as AerK: D216N, T218K, D222K, E254Q and E258Q and the retained positively charged residues highlighted in blue. **B)** I-V curves obtained from the AerK mutant in urea. As the concentration of urea is increased, the open pore current also decreases. **C)** Electrophysiology events from addition of mujdei in cis (1 M KCl, 15 mM HEPES, pH 7.5). Substrate release was very difficult, indicating that the substrate did not translocated the nanopore. **D)** Blank measurement (no substrate added, 1 M KCl, 15 mM HEPES, 2 M urea, pH 7.5). The pore is mostly open, although gating events of variable length, alleviated by bias reversal, do occur in the absence of the substrate. **E)** Addition of the malE219aD10ssrA substrate in cis in the same conditions resulted in substrate entrapment, bias reversal not promoting release and consequently, the pore remained clogged with the substrate. This behaviour is consistent with the one found for the CytK 6K mutant.

A.

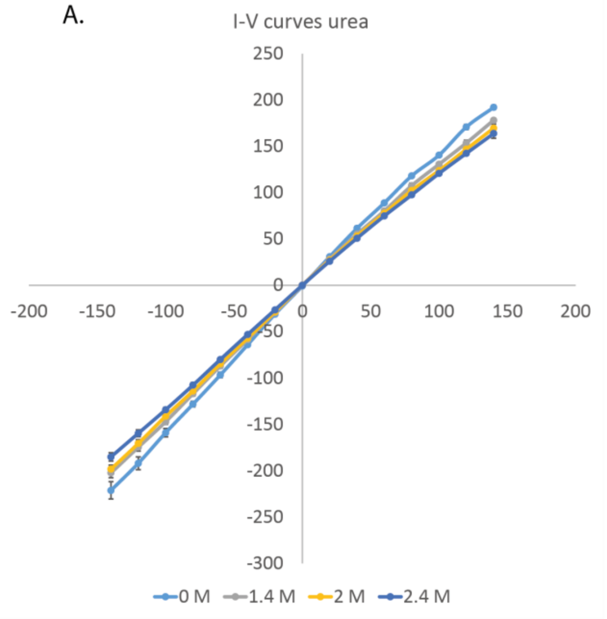

B.

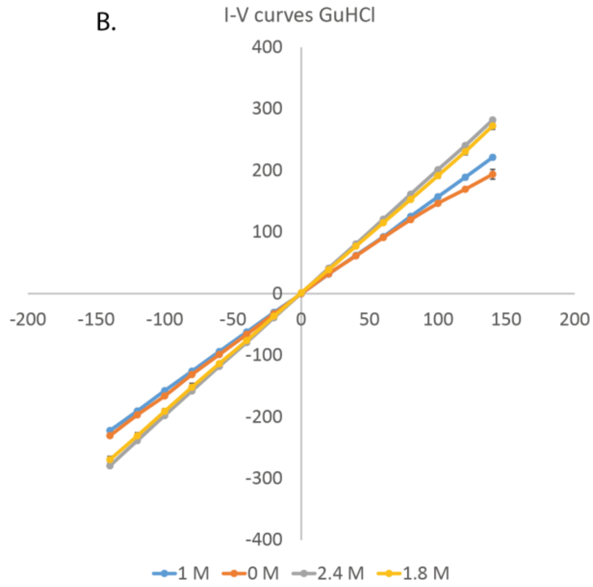

C.

| [urea] (M) | \|I-100\|/ \|I+100\| |
|---|---|
| 0 | 1.13 |
| 1.4 | 1.12 |
| 2 | 1.14 |
| 2.4 | 1.11 |

D.

| [GuHCl] (M) | \|I-100\|/ \|I+100\| |
|---|---|
| 0 | 1.133 |
| 1 | 1.003 |
| 1.8 | 0.998 |
| 2.4 | 0.984 |

Figure **S21. Characterisation of the 2E-4D-CytK nanopore in the two denaturants employed in these experiments. A)** I-V curves from the 2E-4D-CytK nanopore in several concentrations of urea (1.4, 2.0, 2.4 M and no urea). The open pore current decreased with the increase of the urea concentration, while the asymmetry was largely retained. **B)** I-V curves from the 2E-4D-CytK nanopore in several GuHCl concentrations (1.0, 1.8, 2.4 M and no GuHCl). The open pore current increased with the addition of GuHCl, while asymmetry was reversed. **C-D)** The asymmetry of the pore current expressed as the ratio of the absolute value of the current at -100 mV and at +100 mV at the sampled urea (**C**) and GuHCl (**D**) concentrations.

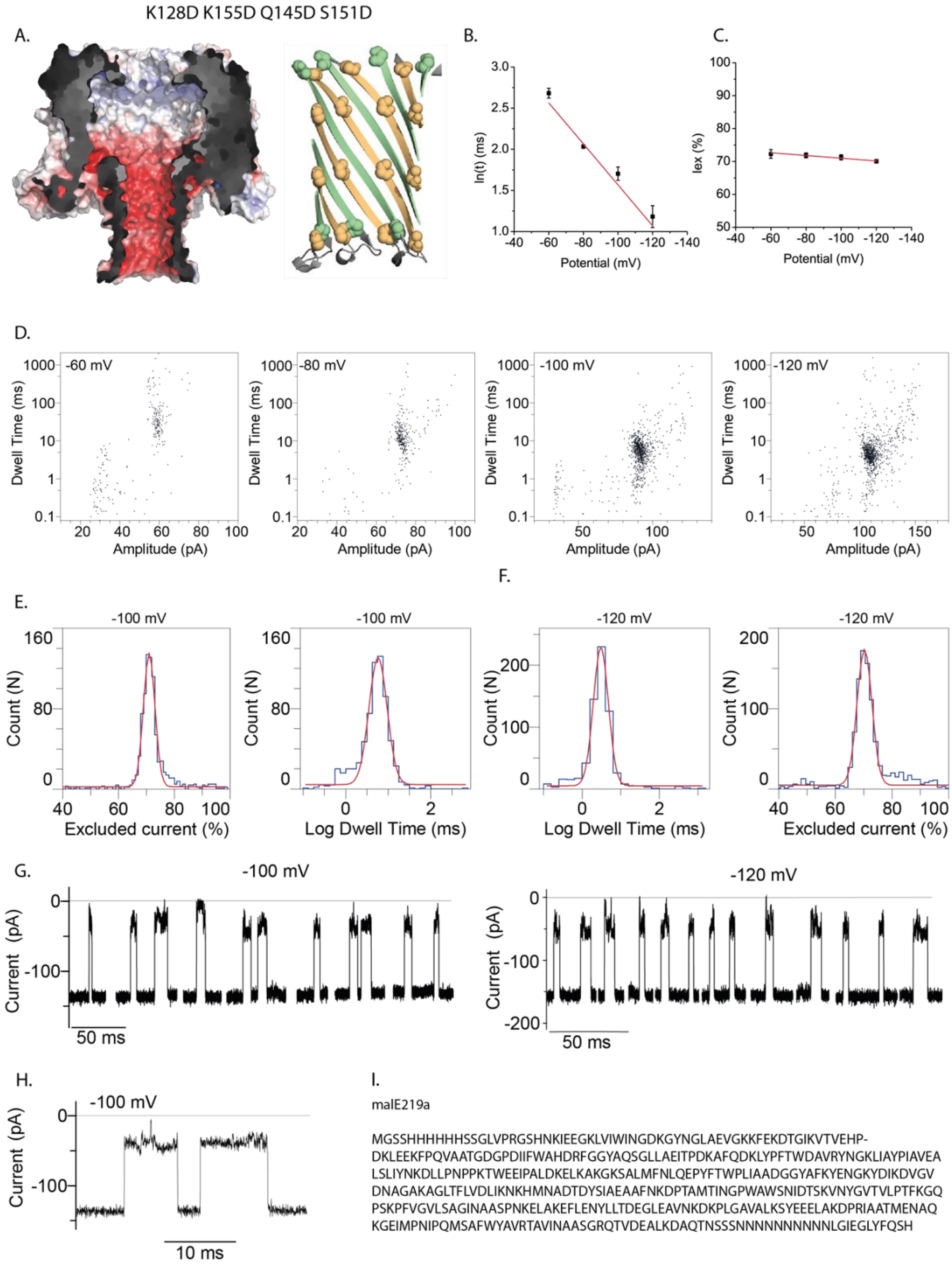

**Figure S22. Translocation of malE219a through CytK-2E-4D nanopores in the presence of 2 M urea. A)** Electrostatic surface illustration of the K128D-K155D-Q145D-S151D-CytK (2E-4D-CytK) nanopore in 1 M KCl at pH 7.5, left; and cartoon representation of the barrel where the substitutions are highlighted, right. **B)** Voltage dependency of the natural logarithm of the dwell time. **C)** $I_{ex\%}$ dependency of malE219a blockades on the applied bias. **D)** Scatter plots (dwell time vs amplitude) associated with malE219a capture and translocation. **E-F)** Examples of histograms obtained for the dwell time and $I_{ex}$ at -100 mV (E) and -120 mV (F). **G)** Typical examples of malE219a capture events at -100 mV (left) and at -120 mV (right). Recordings were carried out in 1 M KCl, 15 mM HEPES, 2 M urea, pH 7.5. **H)** Expanded examples of events at -100 mV. **I)** The protein sequence of malE219a.

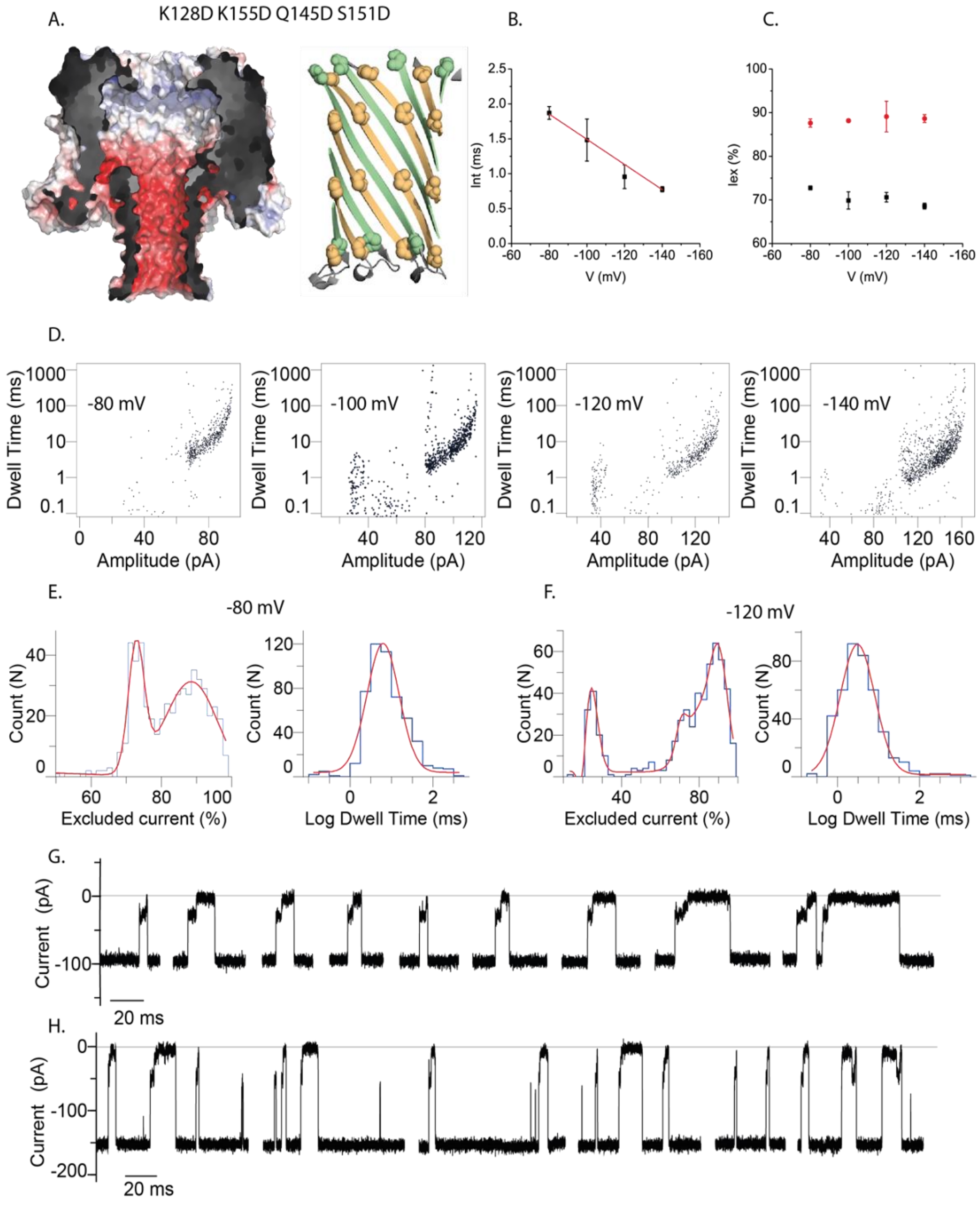

I.

GBP H152A

MANKKVITLSAVMASMLFGAAAHAADTRIGVTIYKYDDNFMSVVRKAIEQDAKAAPDVQLLMNDSQNDQSKQNDQIDVLLAKGVKALAINLVDPAAAGTVIEKAR
GQNVPVVFFNKEPSRKALDSYDKAYYVGTDSKESGIIQGDLIAKHWAANQGWDLNKDGQIQFVLLKGEPGAPDAEARTTYVIKELNDKGIKTEQLQLDTAMWDTAQ
AKDKMDAWLSGPNANKIEVVIANNDAMAMGAVEALKAHNKSSIPVFGVDALPEALALVKSGALAGTVLNDANNQAKATFDLAKNLADGKGAADGTNWKIDNKV
VRVPYVGVDKDNLAEFSKKGSSHHHHH

**Figure S23: Translocation GBP H152A though the 2E-4D CytK nanopore in the presence of 2.4 M urea. A)** Electrostatic surface illustration of the K128D-K155D-Q145D-S151D-CytK (2E-4D-CytK) nanopore in 1 M KCl at pH 7.5, left; and cartoon representation of the barrel where the substitutions are highlighted, right. **B**) Voltage dependency of the natural logarithm of the dwell time. **C)** $I_{ex\%}$ dependency of H152A-GBP blockades on the applied bias. **D)** Scatter plots (dwell time vs amplitude) associated with H152A-GBP capture and translocation. **E-F)** Examples of histograms obtained for the dwell time and $I_{ex}$ at -80 mV (E) and -120 mV (F). **G-H)** Typical examples H152A-GBP capture events at -80 mV (G) and at -120 mV (H). Recordings were carried out in 1 M KCl, 15 mM HEPES, 2.4 M urea, pH 7.5. **I)** The protein sequence of H152A-GBP.

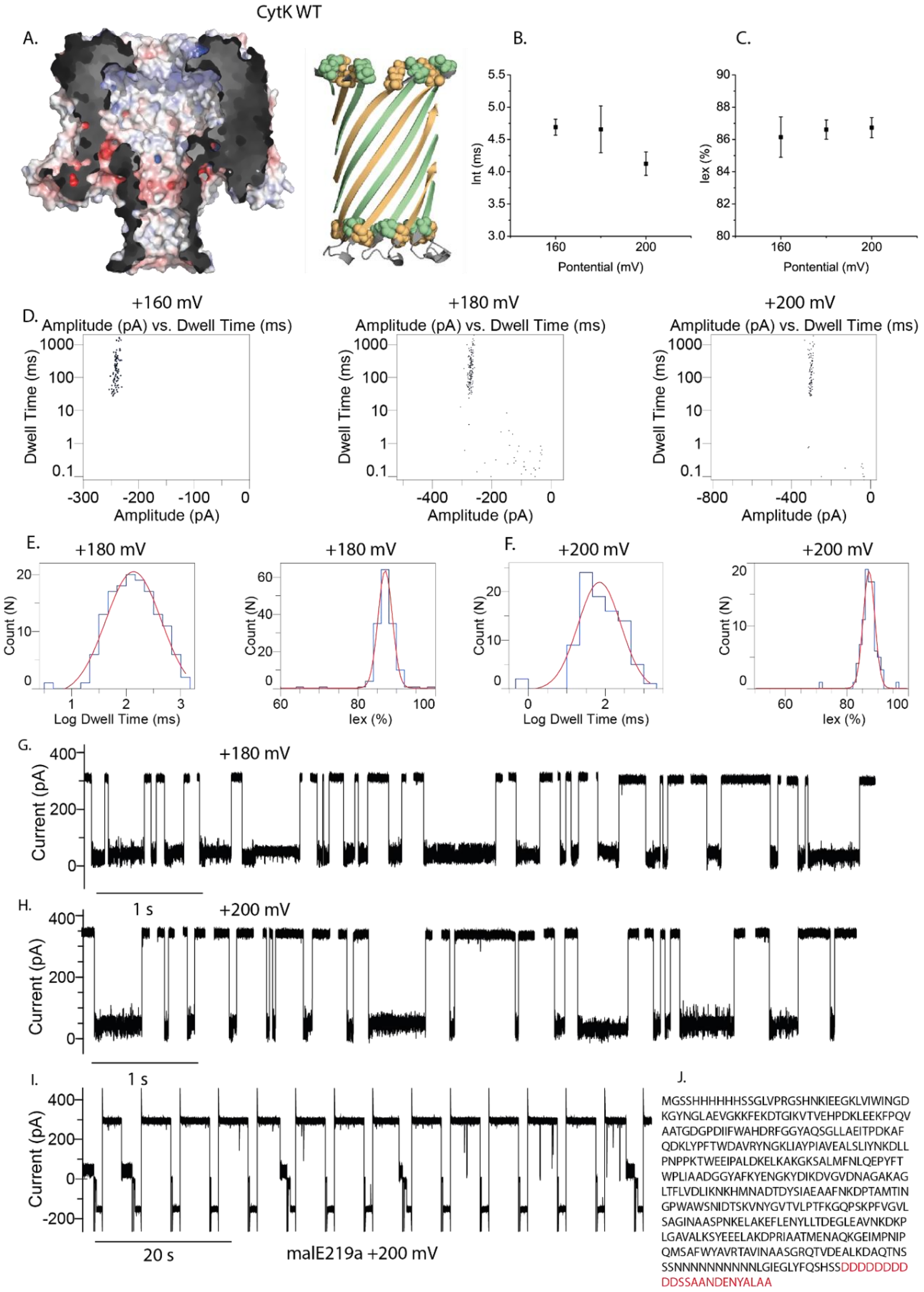

**Figure S24: WY-CytK tested with the malE219a and malE219aD10ssrA proteins in the presence of 1.5 M GuHCl. A)** Electrostatic surface illustration of the WT-CytK nanopore in 1 M KCl at pH 7.5, left; and cartoon representation of the barrel where the substitutions are highlighted, right. **B**) Voltage dependency of the natural logarithm of the dwell time. **C)** $I_{ex\%}$ dependency of malE219a blockades on the applied bias. **D)** Scatter plots (dwell time vs amplitude) associated with malE219a capture and translocation. **E-F)** Examples of histograms obtained for the dwell time and $I_{ex}$ at +180 mV (E) and +200 mV (F). **G-H)** Typical examples of malE219a capture events at +180 mV (G) and at +200 mV (H). Recordings were carried out in 1 M KCl, 15 mM HEPES, 1.5 M GuHCl, pH 7.5. **I)** Capture of malE219a is much more inefficient than for malE219aD10ssrA in the same conditions. **J)** The protein sequence of malE219a (sequence in black) and malE219aD10ssrA (sequence in black followed by sequence in red).

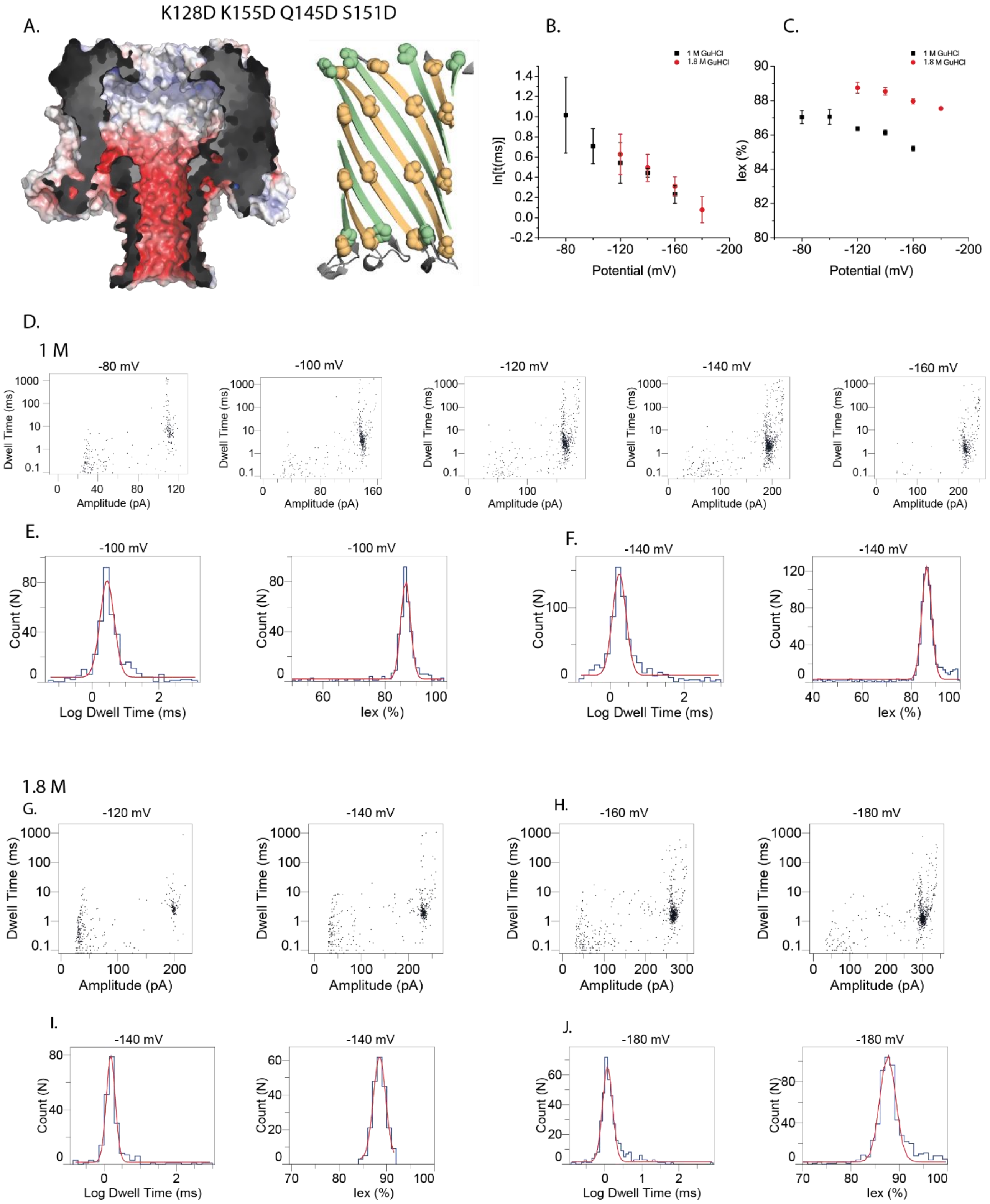
K128D K155D Q145D S151D
A.
B.
C.
1 M GuHCl
1.8 M GuHCl
ln[t(ms)]
Iex (%)
Potential (mV)
D.
1 M
-80 mV
-100 mV
-120 mV
-140 mV
-160 mV
Dwell Time (ms)
Amplitude (pA)
E.
F.
Count (N)
Log Dwell Time (ms)
Iex (%)
1.8 M
G.
H.
I.
J.
-180 mV

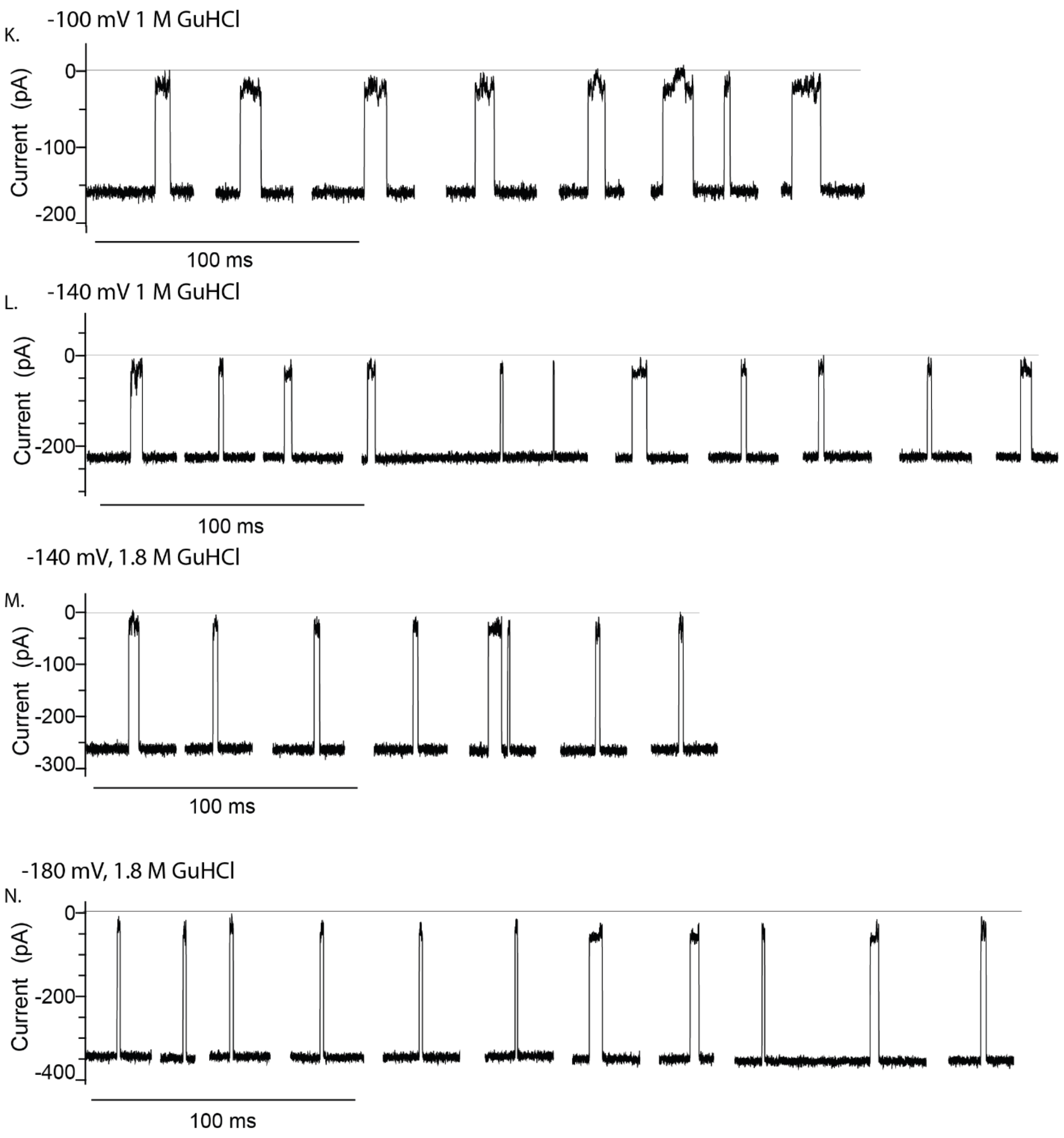

**Figure S25. MalE219a translocation through the 2E-2D CytK mutant in the presence of 1 M and 1.8 M GuHCl. A)** Electrostatic surface illustration of the K128D-K155D-Q145D-S151D-CytK (2E-4D-CytK) nanopore in 1 M KCl at pH 7.5, left; and cartoon representation of the barrel where the substitutions are highlighted, right. **B**) Voltage dependency of the natural logarithm of the dwell time in 1 M (black squares) and 1.8 M (red circles) GuHCl. **C)** $I_{ex\%}$ dependency of malE219a blockades on the applied bias in 1 M (black squares) and 1.8 M (red circles) GuHCl. **D)** Scatter plots (dwell time vs amplitude) associated with malE219a capture and translocation in 1 M GuHCl. **E-F)** Examples of histograms obtained for the dwell time and $I_{ex}$ at -100 mV (E) and -140 mV (F) in 1 M GuHCl. **G-H)** Scatter plots (dwell time vs amplitude) associated with malE219a capture and translocation in 1.8 M GuHCl. **I-J)** Examples of histograms obtained for the dwell time and $I_{ex}$ at -140 mV (I) and -180 mV (J). **K-L)** Typical examples of malE219a capture events at -100 mV (K) and at -140 mV (L). Recordings were carried out in 1 M KCl, 15 mM HEPES, 1 M GuHCl, pH 7.5. **M-N)** Typical examples of malE219a capture events at -140 mV (K) and at -180 mV (L). Recordings were carried out in 1 M KCl, 15 mM HEPES, 1.8 M GuHCl, pH 7.5.

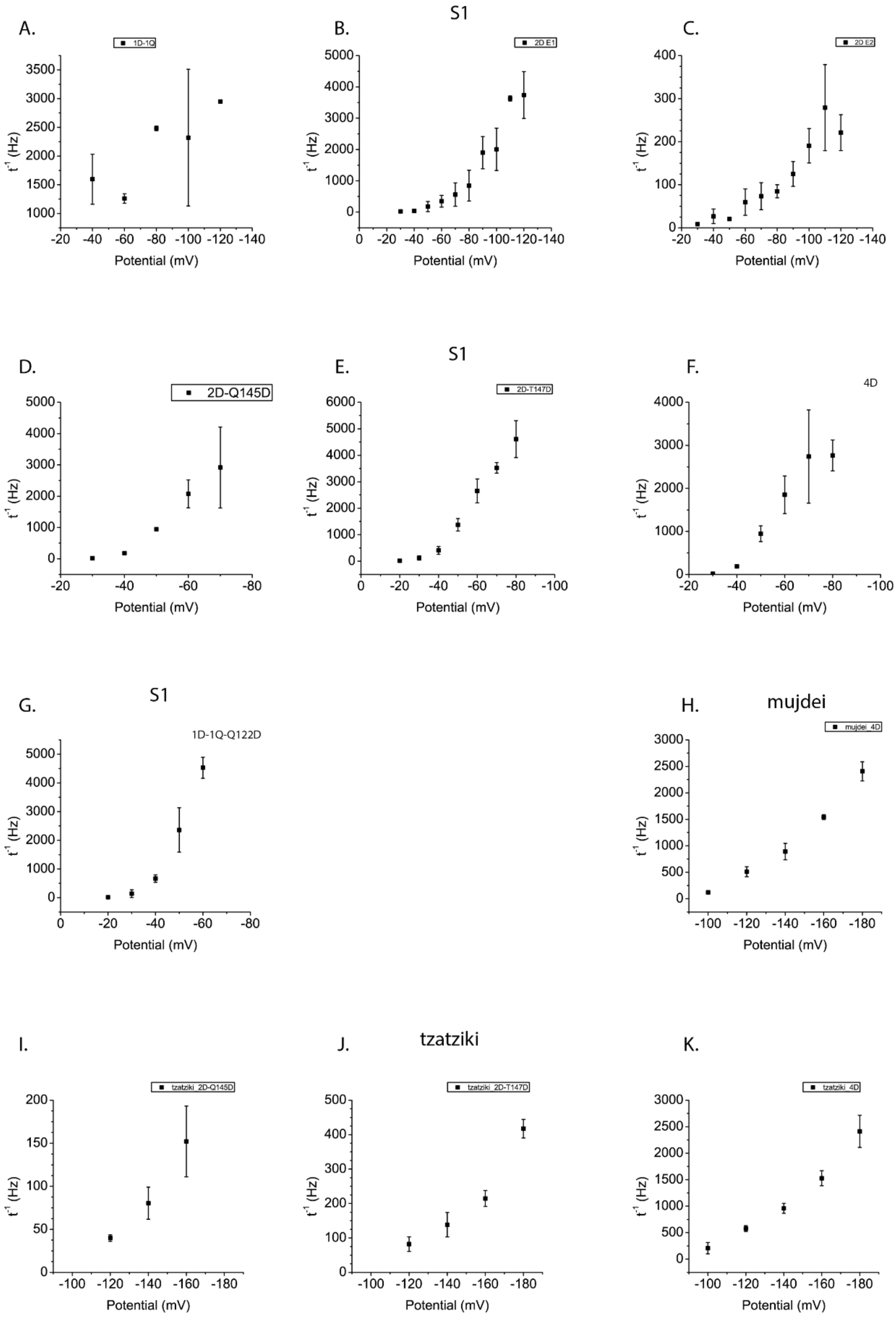

**Figure S26. Translocation rate (1/t) vs the applied potential graphs corresponding to** (**A**) S1 + 2E-1D-1Q-CytK mutant, S1 + 2E-2D-CytK mutant, type 1 events (B) and type 2 events (C), S1 + 2E-2D-Q145D-CytK (D), S1 + 2E-2D-T147D-CytK (E), S1 + 2E-4D-CytK (F), S1 + 2E-1D-1Q-Q122D-CytK (G). The following graphs correspond to the negatively charged model substrates, mujdei + 2E-4D-CytK (H) and tzatziki with 2E-2D-Q145D-CytK (I), 2D-T147D-CytK (J) and 2E-4D-CytK (K).

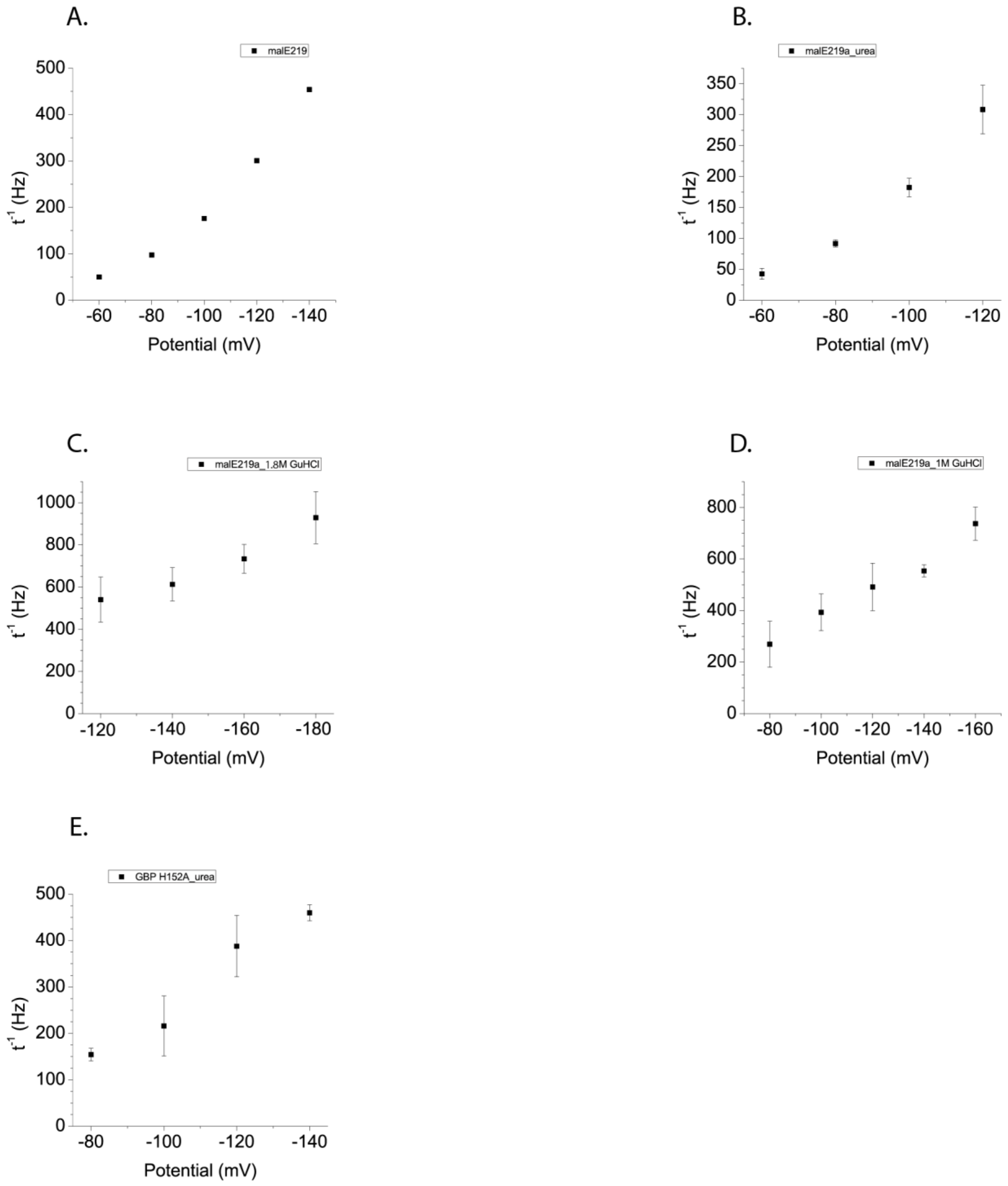

Figure **S27. Rate (1/t) vs the applied bias in the case of the native substrates translocated across the 2E-4D-CytK mutant: malE219 (missing the C-terminal tail) in urea** (A), which gives virtually the same result as malE219a in urea (B), malE219a in 1.8 M GuHCl (C), malE219a in 1 M GuHCl (D) and finally, GBP H152A in urea (E).

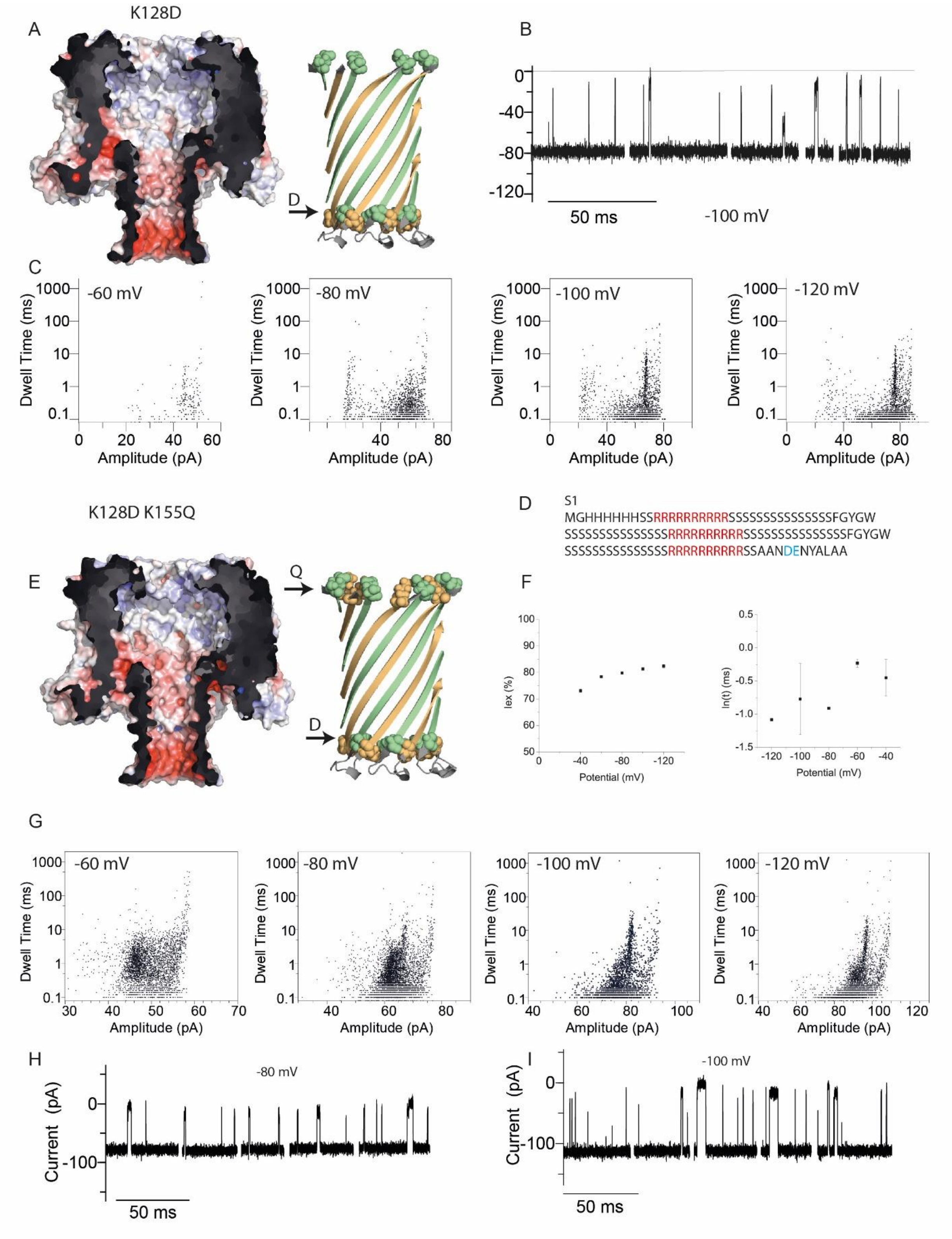

**Figure S28. S1 capture by the K128D-CytK and K128D-K155Q CytK mutants. A)** Electrostatic surface illustration of the K128D-CytK nanopore in 1 M KCl at pH 7.5, left; and cartoon representation of the barrel where the substitutions are highlighted, right. **B)** Typical examples of events of S1 capture by the 2E-1D-CytK nanopore (1 M KCl, 15 mM HEPES, pH 7.5) at -100 mV bias. **C)** Scatter plots (dwell time vs amplitude) associated with S1 capture. **D)** The amino acid sequence of S1, with positively charged residues highlighted in red. **E)** The K128D-K155Q CytK mutant (2E-1D-1Q-CytK) as electrostatic surface representation (1M KCl, pH 7.5), left; and cartoon representation, right. The change of the positively charged Lys to the neutral Gln makes the entry from the vestibule to the barrel weakly more negative than in the WT and the K128D mutant. **F)** The voltage dependencies of the $I_{ex\%}$ and natural logarithm of the dwell time. **G)** Scatter plots obtained electrophysiology traces recorded at several voltages. The removal of the second Lys improved the capture at low potentials, although E1 and E2 are less distinguishable than for the WT. **H-I)** Typical examples of S1 capture events from at -80 mV (H) and at -100 mV (I). Recordings were carried out in 1 M KCl, 15 mM HEPES, pH 7.5.

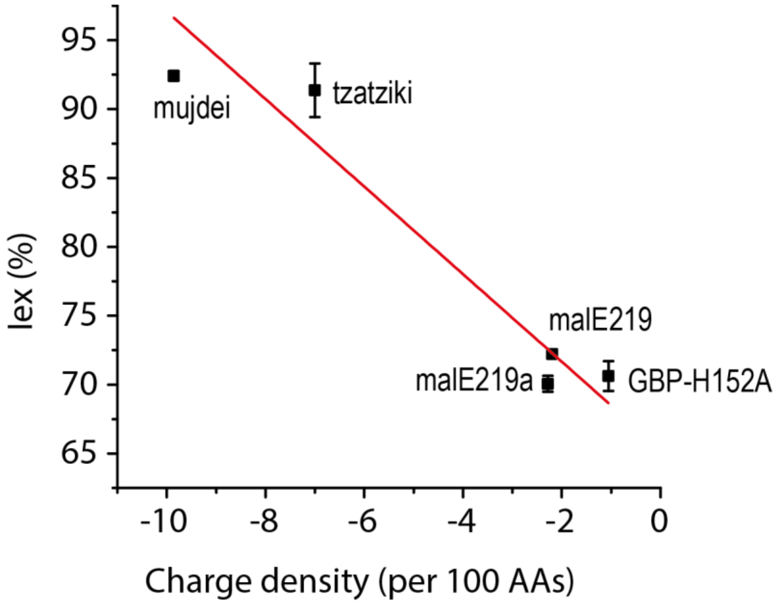

**Figure S29. Variation of the excluded current with the charge density at -120 mV.**

The values of the excluded current at -120 mV, at voltage where all substrates were translocated, were plotted against the charge density of a protein (net charge/length x 100) and an almost linear behaviour emerged. This dependence may be used for fingerprinting proteins. Note: for GBP we used the $I_{ex\%}$ associated with the shallower L1.

The $I_{ex\%}$ may be used in protein identification and potentially in fingerprinting proteins from a mixture. As seen from Figure S25, a somewhat linear dependence was found between the overall excluded current at a voltage where all substrates were translocated and the charge density. Such a relationship may aid protein fingerprinting based on the intrinsic properties of a protein, such as the net charge and length, expressed as charge density. However, for same-charge density proteins, i.e. where the charge density corresponds to the same value, despite the different net charges and lengths, other information may be used to separate these proteins: either analysis of local $I_{ex\%}$ values, or the dwell time and elucidating this extra complexity is beyond the scope of this work.
